# Hypothalamic glucocorticoid receptor in CRF neurons is essential for HPA axis habituation to repeated stressor

**DOI:** 10.1101/2020.11.30.402024

**Authors:** Carine Dournes, Julien Dine, Juan-Pablo Lopez, Elena Brivio, Elmira Anderzhanova, Simone Roeh, Claudia Kuehne, Maria Holzapfel, Rosa-Eva Huettl, Rainer Stoffel, Lisa Tietze, Carola Eggert, Marcel Schieven, Mira Jakovcevski, Jan M. Deussing, Alon Chen

## Abstract

Habituation of the hypothalamic-pituitary-adrenal (HPA) axis to repeated homotypic stressors is crucial for the organism’s well-being. Many physiological and psychological disorders are associated with HPA axis dysfunction. Here, we show that glucocorticoid receptors in CRF neurons of the hypothalamic paraventricular nucleus are essential for HPA habituation. By increasing inhibitory tone onto CRF neurons, glucocorticoid receptors led to essential cellular modulation and hypothalamic-pituitary-adrenal axis activation dampening, when re-exposed to the same stressor.

---

The functionality of the hypothalamic-pituitary-adrenal (HPA) axis is essential for the maintenance of homeostasis and organism’s well-being^1–3^. This includes an appropriate response to a given stressor and its habituation (i.e., a gradual decrease in the magnitude of response to repeated exposure to a relatively low intensity stimulus) following repeated exposure to a homotypic stressor^3^. HPA axis dysfunction has been implicated in the etiology of multiple physiological^1,2,4,5^ and psychological disorders^1,6^. Despite its fundamental role in behavioral and physiological responses to stressful stimuli, the molecular and cellular mechanisms underlying HPA axis habituation are poorly understood.

Parvocellular neurons of the paraventricular nucleus of the hypothalamus (PVN) synthesize and release the neuropeptide corticotropin-releasing factor (CRF)^7^. CRF plays a fundamental and well-established role in the regulation of the HPA axis both under basal and stressful conditions^8,9^. Upon reaching the anterior pituitary gland, CRF stimulates the release of adrenocorticotropic hormone into the general circulation to initiate the synthesis and release of glucocorticoids (GCs; corticosterone (CORT) in rodents and cortisol in humans) from the adrenal cortex^10^. In turn, GCs bind to glucocorticoid and mineralocorticoid receptors (GRs and MRs respectively), which act as transcriptional regulators in the brain and the periphery. GCs also convey the main negative feedback at different levels of the HPA axis^11,12^, particularly onto CRF neurons of the PVN, through GR activation. Thus, PVN-CRF neurons represent the anatomical and functional cells in the brain that mediate the initiation and termination of the neuroendocrine stress response^7^. However, little is known about how GRs regulate the activity of CRF neurons of the PVN, especially in the context of HPA axis habituation. Here, we describe for the first time the role of GRs, expressed specifically by CRF-PVN neurons, in mediating HPA axis habituation to a repeated homotypic stressor. We demonstrate a GR-dependent retrograde signaling mechanism that dampens the activity of CRF neurons of the PVN after exposure to repeated restraint stress, which leads to habituation of the HPA axis.

In order to quantify the colocalization between CRF and GR in neurons of the PVN, we used immunohistochemistry (IHC) on coronal brain sections obtained from *Crf*^*tdTomato*^ (*Crf-ires-Cre*^13^ crossed with Ai9) mice, a well-established CRF reporter mouse line (**Fig. 1a**). We found that 98 ± 0.4 % of CRF^+^ neurons in the PVN co-express GR and conversely 72 ± 1.5 % of GR^+^ cells in the PVN co-express CRF (**Fig. 1a**). Using single-cell RNA (scRNA) sequencing, we profiled the transcriptome of 5,179 cells from the extended PVN. We systematically cataloged their cell type (See **Suppl. Fig. 1a**), under baseline (unstressed) conditions. Our clustering analysis revealed 21 distinct cell populations (10 neuronal and 11 non-neuronal) (**Fig. 1b**, **Suppl. Fig.1b, c and 2**). *Crh* transcript (CRF neurons) was highly expressed in neuronal cluster 3 (**Fig. 1b inset** and **Suppl. Fig. 1b**). Consistent with our immunostaining results, we found that 86% of *Crh*^+^ neurons in cluster 3 also express the *Nr3c1* gene (GR). On the other hand, 49% of *Nr3c1*^+^ neurons co-express *Crh* (**Fig. 1b**). This value is slightly lower than the 72% co-expression observed with IHC (**Fig. 1a**). This is likely due to the dissection method used for the scRNA sequencing. The isolated area is bigger than the actual PVN (**Suppl. Fig. 1a**) and GRs are strongly expressed in this region, leading to the observed “dilution” of the GR/CRF colocalization.

**Figure 1:**
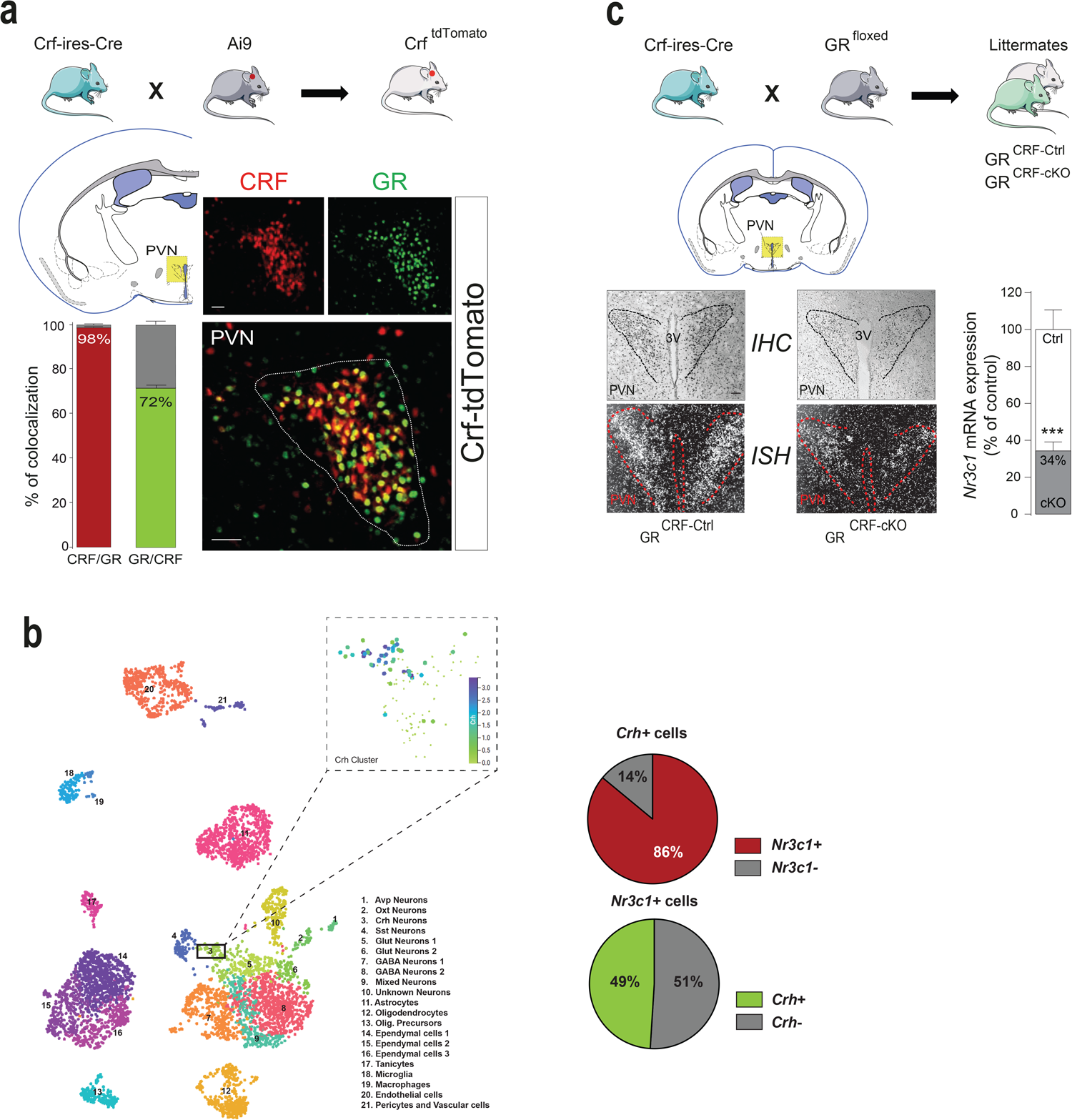
GR expression in CRF^+^ neurons of the PVN and generation of conditional GR deletion in CRF-expressing neurons of the brain. (**a**) Breeding scheme to obtain *Crf*^*tdTomato*^ mice. CRF expression is mapped by tdTomato expression (red) that mirrors the endogenous CRF pattern. Sections are immunostained for glucocorticoid receptor (GR, green channel). 98 ± 0.4% of CRF^+^ neurons of the PVN co-express GR and 72 ± 1.5% of GR^+^ cells co-express CRF (n = 5 mice). Scale bars, 100 μm. (**b**) Uniform Manifold Approximation and Projection (UMAP) plot of 5,179 cells colored per density clustering and annotated according to known cell types. From the neuronal clusters, cluster 3 showed higher expression levels of *Crh* mRNA. 86% of *Crh*^+^ neurons also express *Nr3c1* (red pie chart), while 49% of *Nr3c1*^+^ cells co-express *Crh* (green pie chart). (**c**) Breeding scheme to generate conditional GR knockout mice, *GR*^*CRF-cKO*^, and their control littermates *GR*^*CRF-Ctrl*^. The deletion was validated by assessing GR protein and *Nr3c1* mRNA expression levels using immunohistochemistry (IHC) and *in situ* hybridization (ISH), respectively. *GR*^*CRF-cKO*^ mice express 66 ± 4.8% less *Nr3c1* mRNA in the PVN than *GR*^*CRF-Ctrl*^ mice (unpaired t-test, t_25_ = 6.21, *** p < 0.001, n = 4 control and 4 *GR*^*CRF-cKO*^ mice). PVN, hypothalamic paraventricular nucleus, 3V, third ventricle. Magnification, 20x; scale bar, 100 μm.

To elucidate the physiological role of GRs in CRF^+^ neurons of the PVN, we generated conditional deletion of GRs specifically in CRF^+^ neurons (*GR*^*CRF-cKO*^; **Fig. 1c** and **online methods**) by breeding *Crf-ires-Cre* to floxed GR mice^14^. In *GR*^*CRF-cKO*^ animals, GRs are deleted from all CRF^+^ neurons of the brain, while it is still present in respective control littermates (*GR*^*CRF-Ctrl*^) (**Fig. 1c**). In agreement with our colocalization studies, *GR*^*CRF-cKO*^ animals showed a 66 ± 4.8 % decrease in *Nr3c1* mRNA and GR protein expression in the PVN compared to *GR*^*CRF-Ctrl*^ mice (**Fig. 1c**). CRF/GR colocalization and GR deletion was also observed in the central nucleus of the amygdala (CeA) where CRF^+^ neurons are present. Here, 99 ± 0.4 % of CRF^+^ neurons express GRs but only 28 ± 1.5 % of GR^+^ neurons express CRF (**Suppl. Fig. 3a**). Accordingly, *GR*^*CRF-cKO*^ mice showed a 31 ± 2.6 % decrease in *Nr3c1* mRNA expression in the CeA compared to *GR*^*CRF-Ctrl*^ mice (**Suppl. Fig. 3b**).

Juvenile and adult *GR*^*CRF-cKO*^ mice did not differ in body weight and did not show any gross abnormalities compared to their control littermates (**Suppl. Fig. 3c**). Behavioral characterization revealed no significant differences between *GR*^*CRF-cKO*^ and *GR*^*CRF-Ctrl*^ mice concerning locomotor activity, anxiety-related and stress-coping behaviors (**Suppl. Fig. 4**). To evaluate the consequences of GR deletion from CRF^+^ neurons on circadian rhythmicity of the HPA axis, we measured circulating plasma CORT levels. *GR*^*CRF-cKO*^ mice displayed normal circadian fluctuations in CORT levels with a typical morning trough and afternoon peak. While morning CORT levels were similar in both groups, *GR*^*CRF-cKO*^ mice exhibited an increased plasma CORT peak in the afternoon compared to *GR*^*CRF-Ctrl*^ mice (**Fig. 2a:** [afternoon plasma CORT] in *GR*^*CRF-cKO*^ = 111.60 ± 18.29 ng/ml *vs.* [afternoon plasma CORT] in *GR*^*CRF-Ctrl*^ = 51.65 ± 10.86 ng/ml). We further investigated this difference by continuously measuring free CORT levels within the brain using *in vivo* microdialysis. As with plasma CORT, *GR*^*CRF-cKO*^ mice showed higher CORT levels in the medial prefrontal cortex (mPFC) than *GR*^*CRF-Ctrl*^ animals in the afternoon (**Fig. 2b**).

**Figure 2:**
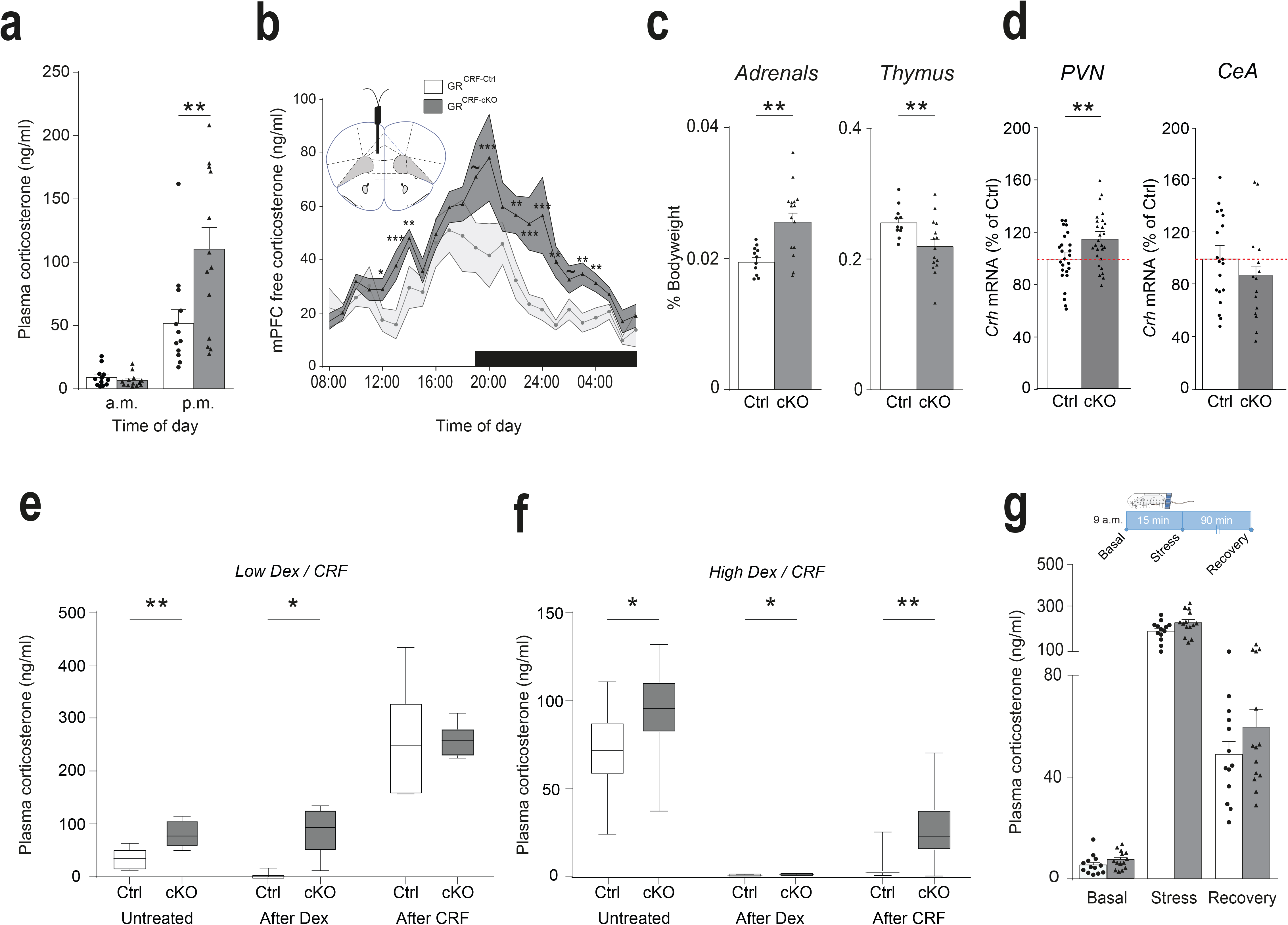
GR deletion in CRF^+^ neurons affects HPA axis rhythmicity, sensitivity and negative feedback, but does not affect HPA axis responsiveness to a single stressor. (**a**) *GR*^*CRF-cKO*^ mice exhibited a higher peak of plasma CORT in the afternoon (p.m.) compared to *GR*^*CRF-Ctrl*^ (2-way RM-ANOVA Genotype * Time of the day F_(1,23)_ = 7.342, p = 0.0125; Bonferroni post-hoc test [afternoon plasma CORT] in *GR*^*CRF-cKO*^ = 111.6 ± 18.29 ng/ml *vs.* [afternoon plasma CORT] in *GR*^*CRF-Ctrl*^ = 51.65 ± 10.8 ng/ml, ** p = 0.0013). n = 12 control and 13 *GR*^*CRF-cKO*^ mice. (**b**) Microdialysis probes were implanted in the mPFC to measure free CORT levels. As for plasma levels, free CORT levels in the mPFC were significantly higher in *GR*^*CRF-cKO*^ mice in the afternoon (2-way RM-ANOVA Time * Genotype F_(2,23)_ = 2.26, p = 0.001; Bonferroni post-hoc test *** p < 0.001, ** p < 0.01, * p < 0.05, ~ p = 0.1). n = 12 control and 13 *GR*^*CRF-cKO*^ mice. (**c**) The adrenal weight (left graph) was significantly higher in *GR*^*CRF-cKO*^ compared to *GR*^*CRF-Ctrl*^ mice (% Bodyweight: 0.020 ± 0.0006 % *vs.* 0.026 ± 0.001 %, unpaired t-test, t_24_ = −3.718, ** p = 0.0011). The thymus weight (right graph) was significantly lower in *GR*^*CRF-cKO*^ compared to *GR*^*CRF-Ctrl*^ mice (% Bodyweight: 0.256 ± 0.007 % *vs.* 0.219 ± 0.011, unpaired t-test, t_24_ = 2.64, ** p = 0.014). n = 11 control and 15 *GR*^*CRF-cKO*^ mice. (**d**) GR deletion in CRF expressing neurons led to increased *Crh* mRNA expression levels under basal conditions in the hypothalamic paraventricular nucleus (PVN) of *GR*^*CRF-cKO*^ mice (left graph; 115.7 ± 4.1% in *GR*^*CRF-cKO*^ mice, unpaired t-test, t_50_ = 2.758, ** p = 0.008; n = 25 control and 27 *GR*^*CRF-cKO*^ mice), but not in the CeA (right graph, 86.7 ± 9.3% in *GR*^*CRF-cKO*^ mice, unpaired t-test, t_32_ = 1.105, p = 0.277; n = 19 control and 15 *GR*^*CRF-cKO*^ mice). (**e-f**) GR deletion in CRF expressing neurons led to a mild increase in HPA axis responsiveness. Plasma CORT levels were measured in response to a pharmacological suppression of adrenocortical activity with either a low (0.05 mg/kg, **e**) or relatively high (2 mg/kg, **f**) dose of Dex and a subsequent stimulation with CRF (0.15 mg/kg). Plasma CORT levels were measured in *GR*^*CRF-Ctrl*^ and *GR*^*CRF-cKO*^ mice 1 week before (Untreated, measure in the afternoon), 6 hours after Dex treatment (After Dex, Dex injection at 9 a.m.) and 30 min after CRF injection (After CRF, CRF injection at 3 p.m.). Data are given as box plots showing medians (lines in the boxes), 25% and 75% percentiles (boxes), as well as the min and max values (whiskers). Statistical differences between the groups are indicated above the columns for each time point (**e**: low dose Dex, unpaired t-test, Untreated: t_18_ = 3.376, p = 0.0034, After Dex: t_18_ = 2.215, p = 0.04, After CRF: t_18_ = 1.096, p = 0.288; **f**: high dose Dex, unpaired t-test, Untreated: t_24_ = 2.299, p = 0.03, After Dex: t_24_ = 2.615, p = 0.015, After CRF: t_24_ = 3.511, p = 0.0018). n = 10 control and 10 *GR*^*CRF-cKO*^ mice in **e** and 10 control and 16 *GR*^*CRF-cKO*^ mice in **f**. (**g**) HPA axis responsiveness to a single restraint stress was not affected by GR deletion. Scheme of the experimental protocol: basal plasma CORT measurement was made at 9 a.m. Plasma CORT levels for the stress response were measured directly after the 15 min of restraint stress and 90 min after the end of the stressor for recovery. No difference was observed between *GR*^*CRF-Ctrl*^ and *GR*^*CRF-cKO*^ mice (2-way RM-ANOVA, Genotype F_(1,13)_ = 3.227, p = 0.096). n = 13 control and 14 *GR*^*CRF-cKO*^ mice.

In accordance with daily higher CORT levels, the adrenal weight was significantly increased in *GR*^*CRF-cKO*^ compared to *GR*^*CRF-Ctrl*^ mice, while thymus weight was reduced (**Fig. 2c**), supporting the observed higher basal CORT levels. As expected, deleting GR from CRF^+^ neurons affected the negative feedback on the HPA axis, which was reflected by the increased levels of *Crh* mRNA in *GR*^*CRF-cKO*^ animals compared to *GR*^*CRF-Ctrl*^ mice in the PVN but not in the CeA (**Fig. 2d**). Since the deletion is specific to CRF^+^ neurons, GR expression levels in the pituitary and the hippocampus of Ctrl and *GR*^*CRF-cKO*^ mice should be similar. qPCR analysis revealed no differences in *Nr3c1* expression levels in the hippocampus (**Suppl. Fig 3d, left**) and surprisingly a slight but significant increase in the pituitary (**Suppl. Fig. 3d, right**), most likely compensating for the daily higher levels of plasma CORT.

Next, HPA axis responsiveness was assessed in *GR*^*CRF-Ctrl*^ and *GR*^*CRF-cKO*^ mice by means of the dexamethasone (Dex)/CRF test (see Methods for details). Two independent tests were performed using either a low or a relatively high dose of Dex to assess GR negative feedback at the level of the pituitary alone or at the level of pituitary and brain, respectively. In both tests, *GR*^*CRF-cKO*^ mice showed significantly higher afternoon CORT levels under basal conditions (**Fig. 2e** and **f**, “Untreated”) compared to *GR*^*CRF-Ctrl*^ mice. This difference is similar to what we previously observed (**Fig. 2a**). As expected, the low dose of Dex strongly suppressed adrenocortical activity in *GR*^*CRF-Ctrl*^ mice, but was ineffective in *GR*^*CRF-cKO*^ (**Fig. 2e**, “After Dex”). Both genotypes responded to stimulation with CRF with a similar increase in CORT levels (**Fig. 2e**, “After CRF”). On the other hand, the high dose of Dex strongly suppressed adrenocortical activity in both genotypes (**Fig. 2f**, “After Dex”). The suppression was maintained in *GR*^*CRF-Ctrl*^ mice even after CRF stimulation, while CRF was able to increase CORT levels in *GR*^*CRF-cKO*^ mice (**Fig. 2f**, “After CRF”). Thus *GR*^*CRF-cKO*^ mice exhibit a weaker negative feedback at the level of the pituitary and a higher sensitivity to CRF than *GR*^*CRF-Ctrl*^ mice.

Finally, to further test the reactivity (and recovery) of the HPA axis in a more physiological paradigm, mice were subjected to a single restraint stress and circulating CORT levels were measured (**Fig. 2g**). Again, plasma CORT levels in the morning, prior to the stressor, were indistinguishable between control and *GR*^*CRF-cKO*^ mice. Surprisingly, no differences in CORT levels were observed immediately following stress exposure nor after recovery (**Fig. 2g**). Taken together, these results demonstrate that deleting GRs from CRF^+^ neurons impacts HPA axis rhythmicity, sensitivity, and negative feedback as expected after releasing the negative brake GRs exert on the HPA axis. Surprisingly however, this deletion does not affect HPA axis responsiveness to an acute single stressor, suggesting GRs are not involved in the regulation of CRF neurons physiology in response to a single stress episode.

To further investigate how GR deletion in CRF^+^ neurons impacts the stress response and especially the habituation of the HPA axis to a repeated homotypic stressor, mice were subjected to a repeated restraint stress protocol (**RRS** see **Fig. 3a** and **online methods**). As shown before, plasma CORT levels in *GR*^*CRF-Ctrl*^ and *GR*^*CRF-cKO*^ mice were indistinguishable under baseline conditions (**Fig. 3b:** solid lines RRS1, RRS2 and RRS3). Surprisingly, upon repetition of the restraint stress, *GR*^*CRF-Ctrl*^ mice exhibited a gradual decrease in plasma CORT levels compared to *GR*^*CRF-cKO*^ animals (dashed orange lines, **Fig. 3b**). This decrease in plasma CORT levels is the result of habituation of the HPA axis response in *GR*^*CRF-Ctrl*^ mice, as shown by the significant difference in CORT level between RRS1 and RRS3 (**Fig. 3b and c**). This habituation is, however, completely absent in *GR*^*CRF-cKO*^ animals (**Fig. 3b and c**). Taken together, these data indicate a role for GRs in CRF neurons during HPA axis habituation.

**Figure 3:**
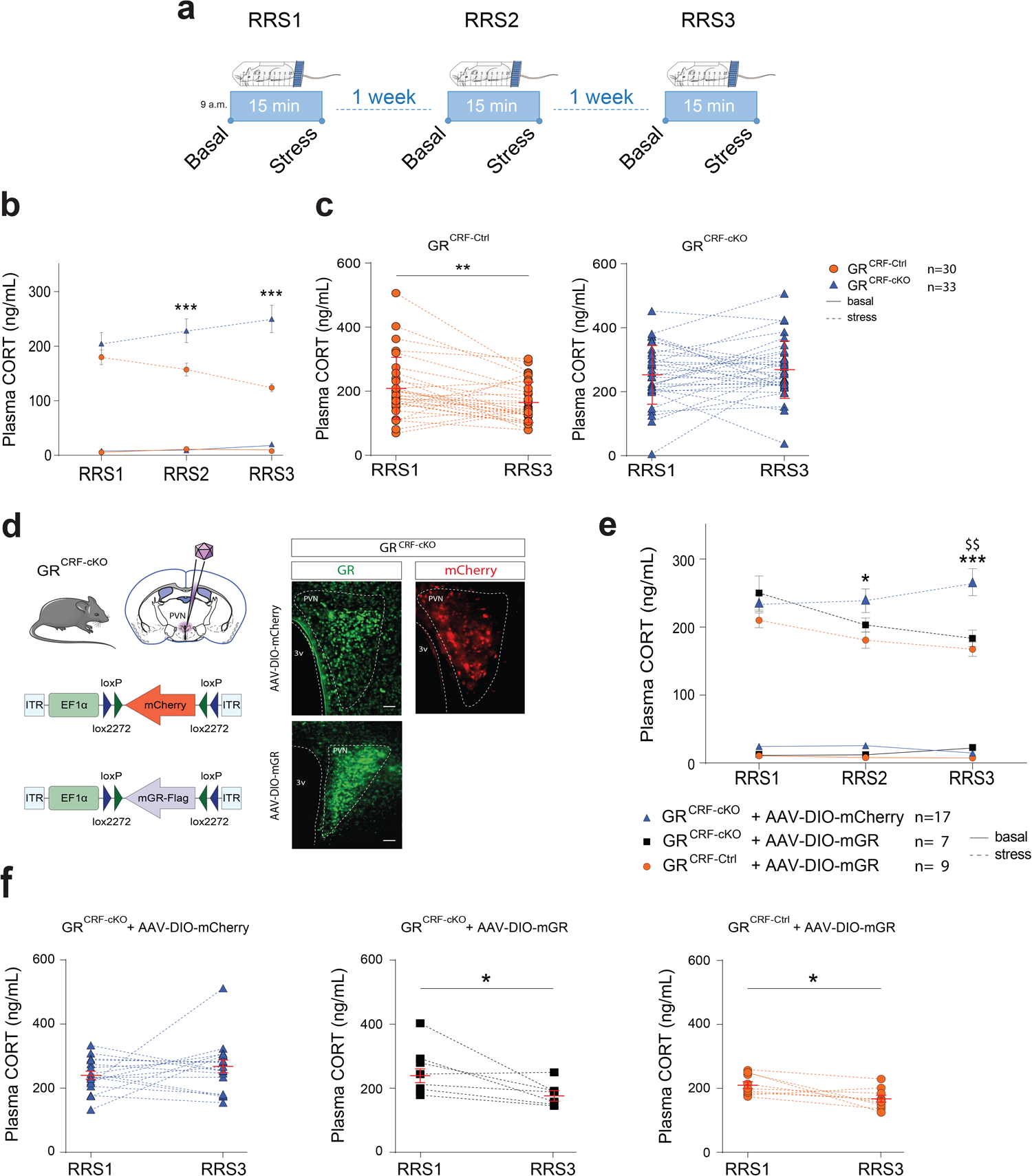
HPA axis habituation to repeated restraint stress (RRS) is mediated by GRs in CRF-expressing neurons of the hypothalamic paraventricular nucleus (PVN). (**a**) Scheme of the Repeated Restraint Stress (RRS) protocol. *GR*^*CRF-Ctrl*^ and *GR*^*CRF-cKO*^ mice were subjected to RRS 3 times with 1-week interval. Corticosterone (CORT) was measured immediately before each stress (“Basal”) and 15 minutes after the end of the stressor (“Stress”). Basal plasma CORT measurement was made at 9 a.m. (**b**) CORT levels were similar under basal conditions (solid lines) between control and *GR*^*CRF-cKO*^ mice for each RRS. However, upon repetition of the RRS, *GR*^*CRF-Ctrl*^ mice showed a progressive decrease in plasma CORT levels after stress compared to *GR*^*CRF-cKO*^ animals (dashed lines: 2-way RM-ANOVA RRS*genotype F_(2,122)_ = 3.36 p = 0.038; Bonferroni post-hoc test *GR*^*CRF-Ctrl*^ *vs. GR*^*CRF-cKO*^ RRS2 and RRS3 *** p< 0.001). (**c**) The decrease in CORT is the result of the habituation of the HPA axis in *GR*^*CRF-Ctrl*^ mice as shown by the direct comparison of plasma CORT levels between RRS1 and RRS3 (left plot, paired t-test *GR*^*CRF-Ctrl*^ RRS1 *vs* RRS3, t_29_ = 3.08, ** p = 0.005, n = 30). This habituation is completely absent in *GR*^*CRF-Ctrl*^ mice (right plot, paired t-test *GR*^*CRF-cKO*^ RRS1 *vs* RRS3, t_32_ = 0.911, p = 0.37, n = 33). (**d**) Design of the viral vector allowing Cre-dependent expression of murine GR: AAV1/2-EF1α-DIO-mGR (AAV-DIO-mGR) or control virus: AAV1/2-EF1α-DIO-mCherry (AAV-DIO-mCherry). The virus was bilaterally delivered into the PVN of *GR*^*CRF-Cre*^ mice (Ctrl = Cre negative and cKO = Cre positive). Four weeks after viral delivery, the correct location of the injection sites was visualized by mCherry expression (AAV-DIO-mCherry) and by the re-expression of the mGR (IHC, green channel) in *GR*^*CRF-cKO*^ mice. Representative 40 μm coronal sections of PVN, scale bars, 100 μm. (**e**) Four weeks after AAV injections mice were subjected to the same RRS protocol showed in (**a**). As before, plasma CORT levels in basal condition were not different between the three groups (solid lines). *GR*^*CRF-Ctrl*^ mice injected with AAV-DIO-mGR showed normal HPA habituation to RRS (dashed line with orange circles) while *GR*^*CRF-cKO*^ mice injected with AAV-DIO-mCherry control virus (dashed line with blue triangles) still lacked habituation. However, the PVN-targeted re-expression of mGR in CRF^+^ neurons in *GR*^*CRF-cKO*^ mice (dashed line with black squares) was sufficient to restore HPA axis habituation to RRS (2-way RM-ANOVA, Genotype effect F_(2,30)_ = 10.97, p = 0.0003; Bonferroni post hoc test *GR*^*CRF-cKO*^ AAV-DIO-mGR *vs. GR*^*CRF-cKO*^ control virus: RRS3 $$ p = 0.005; and *GR*^*CRF-Ctrl*^ AAV-DIO-mGR *vs. GR*^*CRF-cKO*^ AAV-DIO-mCherry control virus: RRS2 * p = 0.041 and RRS3 *** p< 0.001). (**f**) Direct comparison of plasma CORT levels between RRS1 and RRS3 showed the normal habituation of the HPA axis in *GR*^*CRF-Ctrl*^ mice injected with AAV-DIO-mGR virus (right plot: paired t-test, t_8_ = 2.899, p = 0.019, n = 9 mice), and the lack of habituation in *GR*^*CRF-cKO*^ mice injected with AAV-DIO-mCherry control virus (left plot: paired t-test, t_16_ = 1.137, p = 0.27, n = 17 mice). In *GR*^*CRF-cKO*^ mice, the re-expression of mGR only in the PVN is sufficient to restore the HPA axis habituation to RRS (middle plot: paired t-test, t_6_ = 2.672, p = 0.037, n = 7 mice).

To interrogate whether the lack of habituation exclusively depends on the deletion of GR from CRF^+^ neurons of the PVN, we generated a viral vector allowing Cre-dependent expression of murine GR (AAV1/2-Ef1α-DIO-Flag-mGR; AAV-DIO-mGR) (**Fig. 3d**). Bilateral injection of AAV-DIO-mGR into the PVN of *GR*^*CRF-cKO*^ mice led to the specific re-expression of GRs in PVN CRF^+^ neurons (**Fig. 3d**). Four weeks after virus injection, mice were subjected to the same RRS protocol. Again, plasma CORT levels in all three groups of mice were indistinguishable under baseline conditions (**Fig. 3e:** solid lines RRS1, RRS2 and RRS3). *GR*^*CRF-Ctrl*^ mice injected with AAV-DIO-mGR (**Fig. 3e** dotted line with orange circles, and **f** right plot) showed a normal habituation to RRS, comparable to the one observed in non-injected *GR*^*CRF-Ctrl*^ mice (**Fig 3b**). In contrast, *GR*^*CRF-cKO*^ mice injected with control AAV-DIO-mCherry confirmed the previously observed lack of habituation of the HPA axis (**Fig. 3e** dotted line with blue triangles and **f** left plot). Notably, the re-expression of GRs only in the PVN of *GR*^*CRF-cKO*^ mice was sufficient to restore the habituation of the HPA axis to the RRS (**Fig. 3e** dotted line with black squares and **f** middle plot).

We next investigated the possible mechanisms underlying this GR-dependent habituation of the HPA axis to RRS in *GR*^*CRF-Ctrl*^ mice. After RRS, *GR*^*CRF-cKO*^ showed higher *Crh* mRNA expression in the PVN compared to *GR*^*CRF-Ctrl*^ mice, as well as higher levels of stress-induced *cFos* mRNA (**Fig. 4a**), suggesting higher stress-induced CRF expression and neuronal activity in the PVN, thus leading to higher plasma CORT levels. Thus, we hypothesized that a change in intrinsic properties and/or excitability of CRF^+^ neurons of the PVN could lead to the decreased CORT release observed after RRS in *GR*^*CRF-Ctrl*^ mice^12^. Targeted patch-clamp recordings of CRF-tdTomato neurons in the PVN (**Fig. 4b**) showed no genotype or stress effect on intrinsic electrical properties (resting membrane potential [RMP], input resistance) and excitability of these cells (**Fig. 4c,** representative traces and **4d, e, f,** for RMP, input resistance and excitability, respectively). However, RRS induced a decrease in miniature excitatory post-synaptic currents (mEPSCs) frequency (**Fig. 4g and h**: *GR*^*CRF-Ctrl*^ Basal *vs.* RRS: 0.75 ± 0.07 Hz *vs.* 0.47 ± 0.05 Hz) and an increase in miniature inhibitory post-synaptic currents (mIPSCs) frequency (**Fig. 4i and j**: *GR*^*CRF-Ctrl*^ Basal *vs.* RRS: 8.42 ± 0.94 Hz *vs.* 17.50 ± 1.51 Hz) in *GR*^*CRF-Ctrl*^ mice without affecting the amplitude of the miniature currents (**Suppl. Fig. 5a and b** for mEPSCs and mIPSCs respectively), suggesting a presynaptic modification in the release probability of neurotransmitters (decreased for glutamate and increased for GABA). These RRS-induced effects were absent in *GR*^*CRF-cKO*^ mice (**Fig. 4g**: mEPSCs frequency: *GR*^*CRF-cKO*^ Basal *vs.* RRS: 0.83 ± 0.06 Hz *vs.* 0.80 ± 0.08 Hz; **Fig. 4i**: mIPSCs frequency: *GR*^*CRF-cKO*^ Basal *vs.* RRS: 9.38 ± 1.18 Hz *vs.* 10.69 ± 0.98 Hz). Taken together, our results demonstrate that HPA axis habituation to RRS is mediated by a GR-dependent switch in the excitatory / inhibitory synaptic transmission onto CRF^+^ neurons of the PVN.

**Figure 4:**
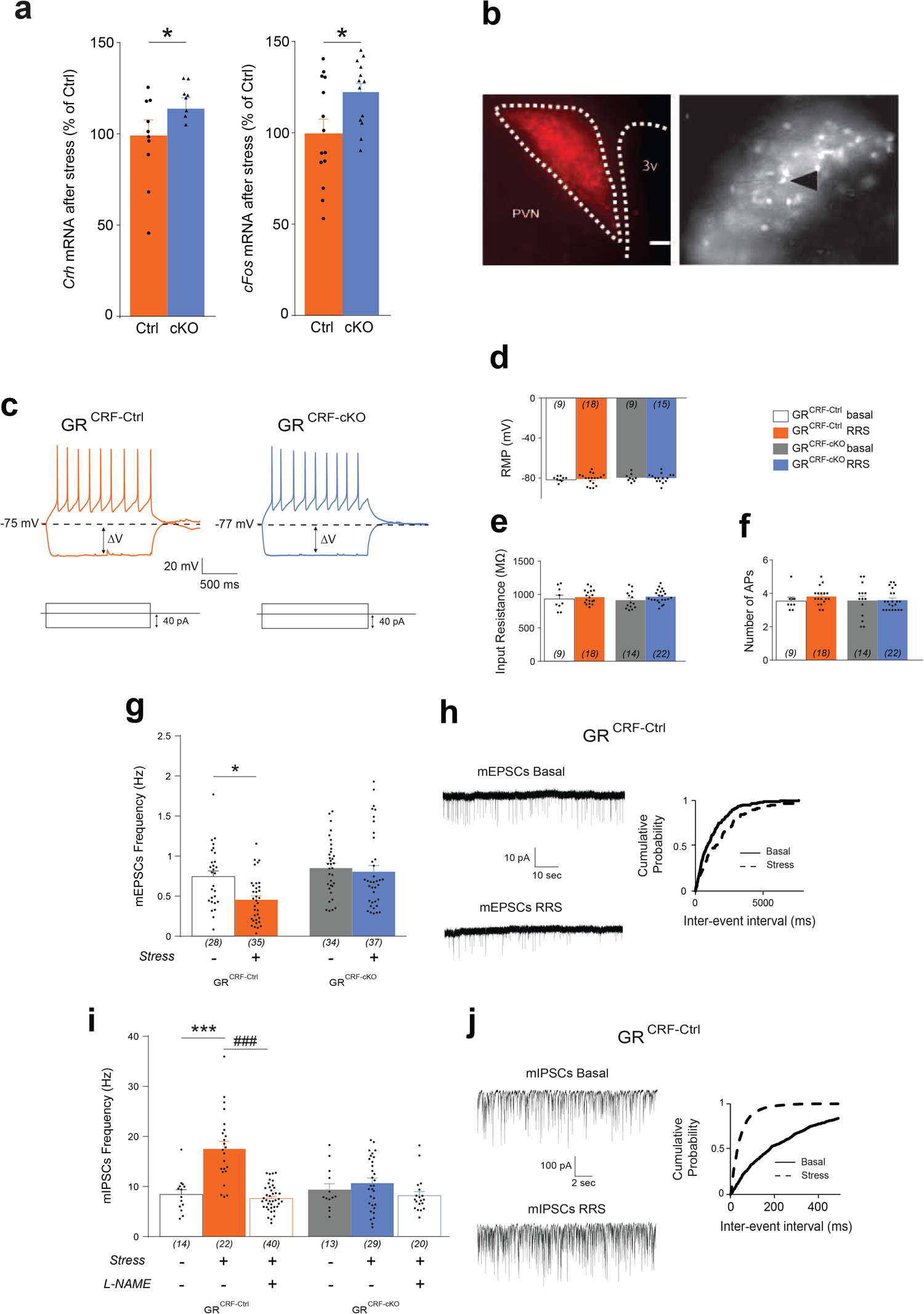
GR deletion in CRF^+^ neurons impairs the stress-induced excitation / inhibition switch responsible for the habituation of the HPA axis. (**a**) Upon repeated restraint stress (RRS), *GR*^*CRF-cKO*^ mice showed higher *Crh* mRNA expression in the PVN (left bar graph: unpaired t-test, t_16_ = 2.429, p = 0.02; n = 10 control and 8 cKO mice), as well as *cFos* mRNA (right bar graph: unpaired t-test, t_25_ = 2.371, p = 0.026; n = 14 control and 13 *GR*^*CRF-cKO*^ mice) compared to *GR*^*CRF-Ctrl*^. (**b**) Microphotograph from a 300 μm coronal acute brain slice of the PVN at 5x (left) and 40x (right) magnification. CRF neurons of the PVN were visually identified by the expression of tdTomato (black arrow shows an example of a recorded cell). PVN, hypothalamic paraventricular nucleus, 3V, third ventricle. Scale bar, 100 μm. (**c**) Representative traces from CRF neurons recordings (current-clamp mode) from *GR*^*CRF-Ctrl*^ (orange) and *GR*^*CRF-cKO*^ (blue) mice. (**d-f**) The intrinsic membrane properties of CRF neurons from Ctrl and *GR*^*CRF-cKO*^ mice recorded under basal and stress (RRS protocol) conditions were not different (2-way ANOVAs): resting membrane potential (RMP, **d**), input resistance (**e**), and excitability (**f**). Cell numbers are indicated in brackets in each bar and were obtained from at least 7 mice. (**g**) RRS induced a strong decrease in miniature excitatory post-synaptic currents (mEPSCs) frequency in *GR*^*CRF-Ctrl*^ but not in *GR*^*CRF-cKO*^ mice (2-way ANOVA Stress effect F_(1,130)_ = 6.335, p = 0.013, Bonferroni post-hoc test *GR*^*CRF-Ctrl*^ Basal *vs.* RRS: 0.75 ± 0.07 Hz *vs*. 0.47 ± 0.05 Hz, p = 0.02; *GR*^*CRF-cKO*^ Basal *vs*. RRS: 0.83 ± 0.06 Hz *vs.* 0.80 ± 0.08 Hz, p = 0.9). Cell numbers are indicated in brackets under each bar and were obtained from at least 7 mice. (**h**) Representative traces and cumulative plot of inter-event interval distribution of mEPSCs in *GR*^*CRF-Ctrl*^ mice under basal (black line) and stress (dotted line) conditions. Stress induced a rightward shift of the distribution reflecting a decrease in mEPSCs frequency. (**i**) RRS induced a strong increase in miniature inhibitory post-synaptic currents (mIPSCs) frequency in *GR*^*CRF-Ctrl*^ but not in *GR*^*CRF-cKO*^ mice (2-way ANOVA Stress*Genotype F_(1,74)_ = 8.76, p = 0.004, Bonferroni post-hoc test *GR*^*CRF-Ctrl*^ Basal *vs.* RRS: 8.42 ± 0.94 Hz *vs.* 17.50 ± 1.51 Hz, *** p < 0.001; *GR*^*CRF-cKO*^ Basal *vs.* RRS: 9.38 ± 1.18 Hz *vs.* 10.69 ± 0.98 Hz, p = 0.9). Moreover, the incubation of PVN slices from *GR*^*CRF-Ctrl*^ mice in the nitric oxide synthase 1 inhibitor L-NAME was able to completely reverse this RRS-induced increase in mIPSCs frequency (*GR*^*CRF-Ctrl*^ RRS = 17.50 ± 1.51 Hz *vs. GR*^*CRF-Ctrl*^ RRS + L-NAME = 7.63 ± 0.41 Hz, unpaired t-test, t_60_ = 7.92, ### p < 0.001), without affecting the mIPSCs frequency in *GR*^*CRF-cKO*^ mice (*GR*^*CRF-cKO*^ RRS = 10.69 ± 0.98 Hz *vs. GR*^*CRF-cKO*^ RRS + L-NAME = 8.19 ± 0.78 Hz, unpaired t-test, t_47_ = 1.84, p = 0.072). Cell numbers are indicated in brackets under each bar and were obtained from at least 7 mice. (**j**) Representative traces and cumulative plot of inter-event interval distribution of mIPSCs in *GR*^*CRF-Ctrl*^ mice under basal (black line) and stress (dotted line) conditions. Stress induced a leftward shift of the distribution reflecting an increase in mIPSCs frequency.

GRs are expressed post-synaptically but modulate synaptic transmission in the context of RRS. This suggests the involvement of a retrograde messenger, which is released from the post-synapse in a GR-dependent manner to act at the pre-synaptic level to change the release probability of neurotransmitters^15^. Both the endocannabinoid (eCB) and nitric oxide (NO) systems are very potent retrograde modulators of synaptic transmission in the PVN^16–19^. Accordingly, the RRS-induced increased inhibition observed in *GR*^*CRF-Ctrl*^ mice was completely reversed by the inhibition of the nitric oxide synthase 1 (NOS1) by pre-incubating the slices in L-NAME (100 μM) (**Fig. 4i**: mIPSCs frequency: *GR*^*CRF-CtrlRRS*^ *vs.* RRS + L-NAME: 17.50 ± 1.51 Hz *vs.* 7.63 ± 0.41 Hz), again without affecting the amplitude of mIPSCs (**Suppl. Fig. 5b**), nor exerting any effect in *GR*^*CRF-cKO*^ RRS mice. This suggests an RRS-mediated GR-dependent release of NO, which acts retrogradely at the presynaptic terminal to increase the release probability of GABA onto CRF^+^ neurons of the PVN.

To summarize, GR deletion from CRF^+^ neurons does not affect their intrinsic electrical properties nor the HPA axis response to a single stressor. However, we showed that, upon repetition of the same stressor (RRS), control mice exhibit habituation of the HPA axis, as shown by reduced plasma CORT levels. The HPA axis habituation is dependent on GR expression in CRF neurons of the PVN and does not involve changes in intrinsic electrical properties of these neurons. These findings contrast with a recently published study showing CORT-independent fast adaptation of CRF neurons excitability to repeated white-noise presentation, measured by *in vivo* calcium imaging^20^. In our study, the HPA habituation is likely mediated by a change in the excitation / inhibition balance, towards a stronger inhibitory tone onto CRF^+^ neurons of the PVN, leading to lower CORT release from the adrenals. Although the mechanism underlying the decreased excitation has yet to be determined, we showed that the increased inhibition is mediated by GR-dependent retrograde action of NO at the presynaptic side, leading to an increased release probability of GABA. Previous studies in mice and rats have demonstrated that the eCB and the NO systems are very potent neuromodulators of synaptic transmission onto magnocellular and parvocellular neurons of the PVN^16–19^. Hence, repetitive immobilization induced a genomic GR-dependent impairment of eCB signaling at GABAergic and glutamatergic synapses on parvocellular PVN neurons^16^, however, without discriminating between the different populations of parvocellular neurons. Other studies showed a rapid non-genomic and synapse-specific GC effect driven by retrograde eCB signaling at glutamatergic synapses and by NO signaling at GABAergic synapses on magnocellular and parvocellular PVN neurons^17–19^. This modulation leads to strong changes in the electrical activity of these neurons and thus in peptide release (i.e. oxytocin, vasopressin, thyrotropin-releasing hormone, and CRF). Moreover, it was demonstrated that the rapid non-genomic GC effect requires expression of the classical nuclear GR, since its deletion in the PVN completely abolishes the effect on synaptic transmission^17^. It is nonetheless unclear whether this mechanism takes place endogenously. Since Dex was bath-applied on brain slices, it is difficult to draw definitive conclusions about the physiological regulation of neuronal networks underlying stress responses. Here, we showed that a similar GR-dependent retrograde signaling is an endogenous key component of HPA axis habituation to a repeated stressor by dampening synaptic transmission onto CRF^+^ neurons of the PVN.

## Online methods

Methods, including statements of data availability and any associated accession codes and references, are available in the online version of the paper.

## Acknowledgments

We thank C. Wotjak, M. Schmidt, G. Balsevich and M. Eder for valuable discussions on the project. We thank A. Varga and all the caretakers of the Max Planck Institute of Psychiatry for their devoted assistance with animal care. We thank J. Keverne for professional English editing, and S. Karamihalev for help with graphical design. We thank T. Rein for the guide plasmid pRK5-mGR-Flag.

A.C. is the incumbent of the Vera and John Schwartz Family Professorial Chair at the Weizmann Institute and is the head of the Max Planck Society–Weizmann Institute of Science Laboratory for Experimental Neuropsychiatry and Behavioral Neurogenetics. This work is supported by: an FP7 Grant from the European Research Council (260463, A.C.); a research grant from the Israel Science Foundation (1565/15, A.C.); the ERANET Program, supported by the Chief Scientist Office of the Israeli Ministry of Health (3-11389, A.C.); the project was funded by the Federal Ministry of Education and Research under the funding code 01KU1501A (A.C.); research support from Roberto and Renata Ruhman (A.C.); research support from Bruno and Simone Licht; I-CORE Program of the Planning and Budgeting Committee and The Israel Science Foundation (grant no. 1916/12 to A.C.); the Nella and Leon Benoziyo Center for Neurological Diseases (A.C.); the Henry Chanoch Krenter Institute for Biomedical Imaging and Genomics (A.C.); the Perlman Family Foundation, founded by Louis L. and Anita M. Perlman (A.C.); the Adelis Foundation (A.C.); the Marc Besen and the Pratt Foundation (A.C.); and the Irving I. Moskowitz Foundation (A.C.).

J.D. held a Max Planck Society Fellowship and is now supported by the Achar family as the incumbent of the Achar Research Fellow Chair in Electrophysiology of the Weizmann Institute of Science.

J.M.D. is supported by the Max Planck Society and part of the work was supported by the German Federal Ministry of Education and Research (FKZ:01ZX1314H).

J.P.L. holds postdoctoral fellowships from the European Molecular Biology Organization (EMBO-ALTF 650-2016), Alexander von Humboldt Foundation, and the Canadian Biomarker Integration Network in Depression (CAN-BIND).

M.J. held a NARSAD Young Investigator Grant #22803.

## Author contributions

C.D., J.D., J-P.L., E.B., E.A., C.K., and M.J., designed and performed the experiments. J.D. designed and performed electrophysiological studies. S.R. performed bioinformatics analysis. M.H., R-E.H, R.S., L.T., and C.E. assisted in experiments. C.D., J.D., J.M.D., and A.C conceived, designed and supervised the project. J.D, C.D., J-M.D. and A.C. wrote the manuscript. All authors revised and approved the final version of the manuscript to be published.

---

**Supplementary figure 1:**
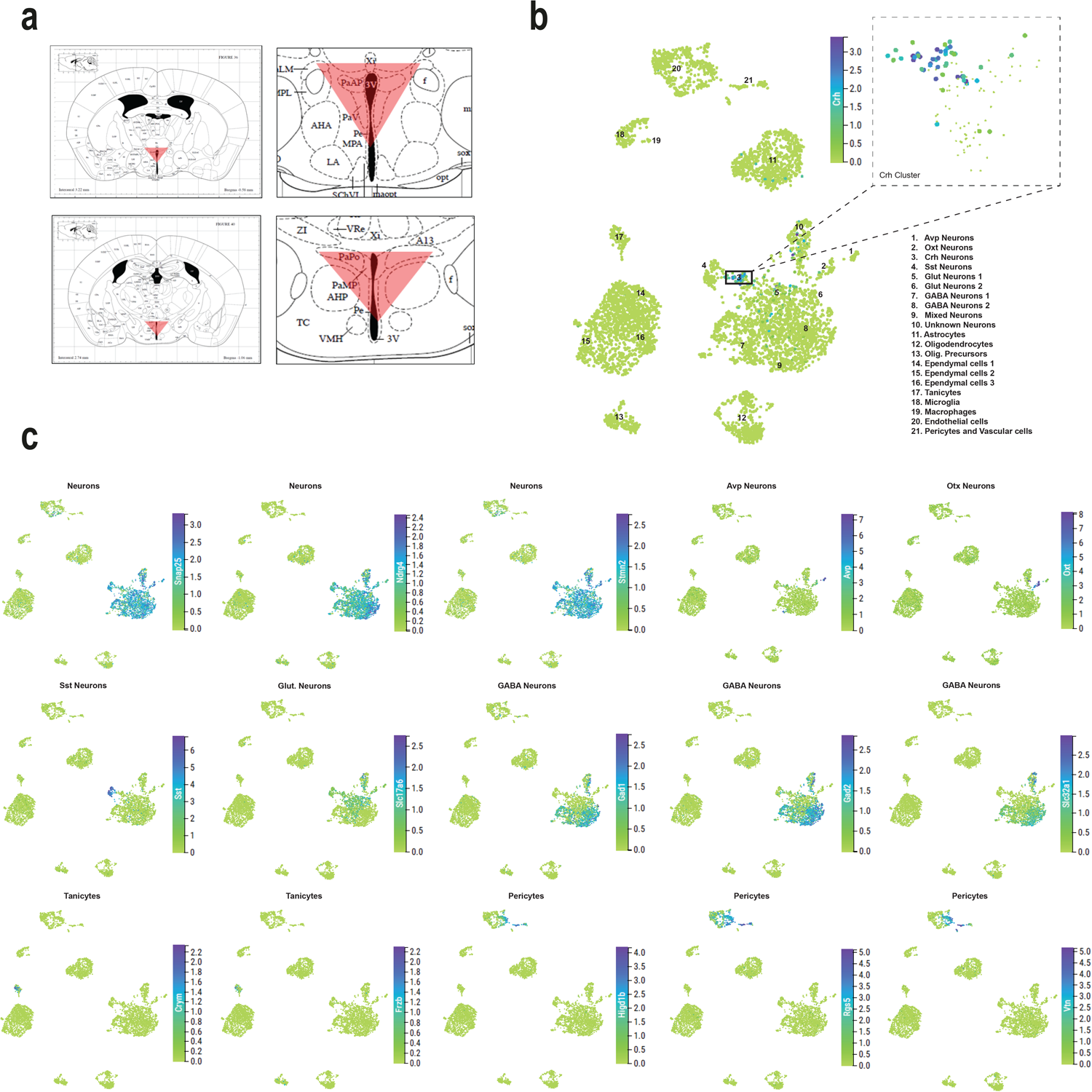
Transcriptomic profile of the mouse hypothalamic paraventricular nucleus (PVN) and *Crh* expression. (**a**) Scheme depicting the dissection of the PVN for single-cell RNA (scRNA) sequencing experiment (adapted from the Paxinos mouse brain atlas). (**b**) UMAP plot of 5,179 cells colored per density clustering and annotated according to known cell types. While *Crh* mRNA is weakly expressed in non-neuronal cell populations (except few astrocytes), its expression is higher in neuronal cells and especially highly enriched in cluster 3 (*Crh* cluster). (**c**) UMAP plots showing expression levels of the marker genes for each cluster identified.

**Supplementary figure 2:**
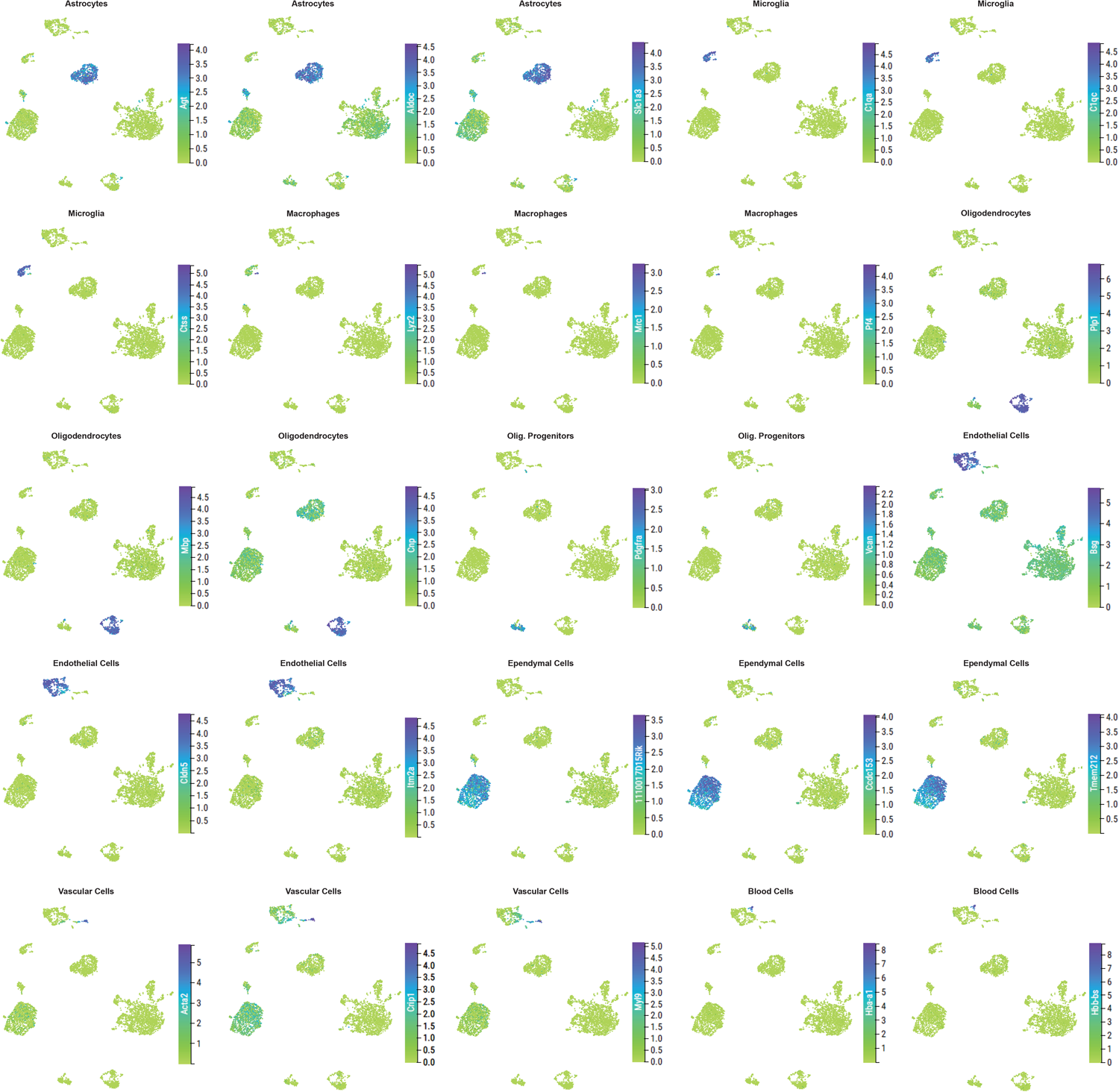
Transcriptomic profile of the mouse hypothalamic paraventricular nucleus (PVN). UMAP plots showing expression levels of the marker genes for each cluster identified

**Supplementary figure 3:**
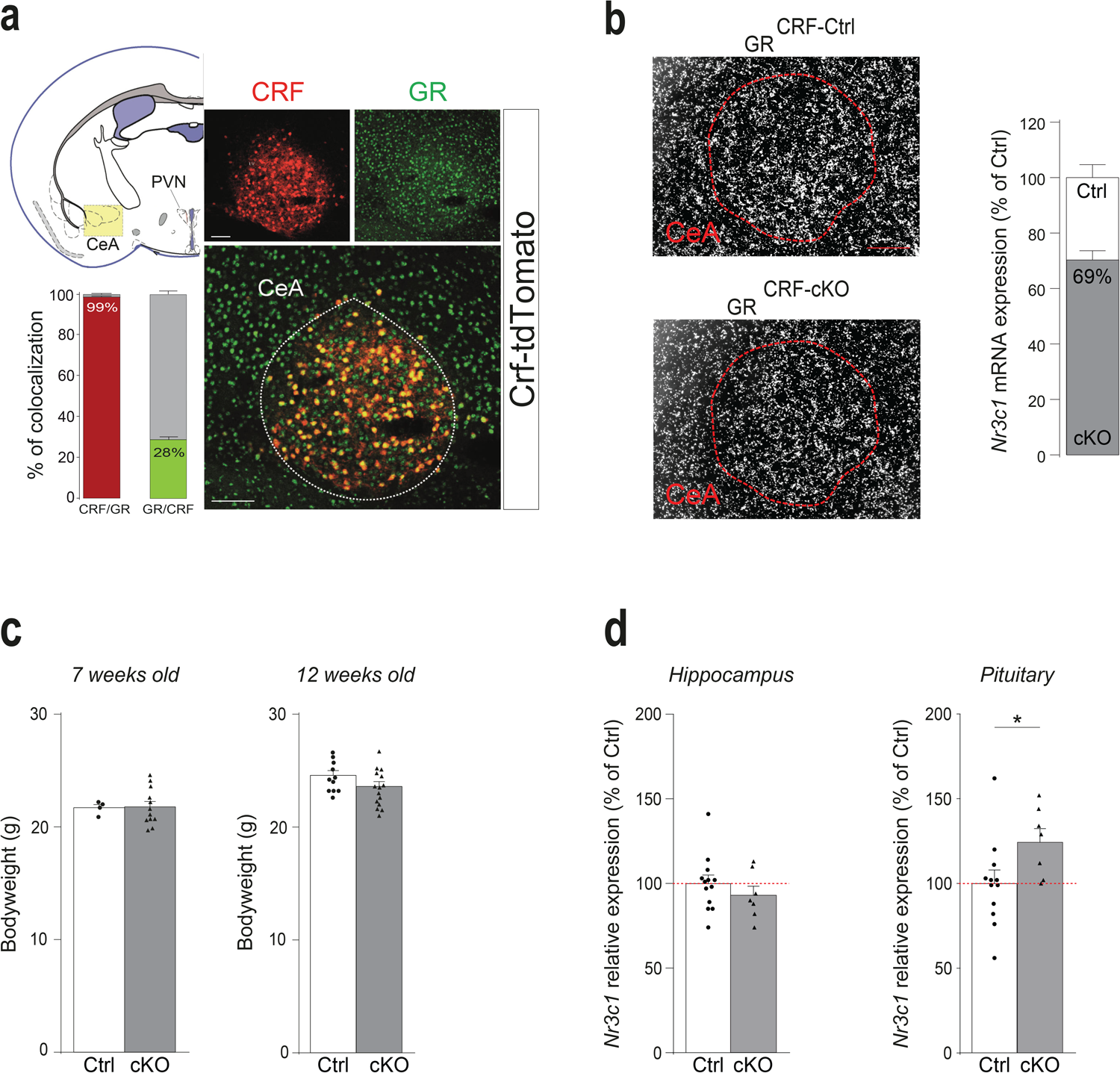
Characterization of GR deletion in CRF expressing neurons. (**a**) CRF / GR colocalization was also observed in the central nucleus of the amygdala (CeA). We found that 99 ± 0.4% of CRF^+^ neurons express GR but only 28 ± 1.5% of GR^+^ neurons co-express CRF (n = 5 mice). Scale bars, 50 μm. (**b**) *GR*^*CRF-cKO*^ mice showed a 31 ± 2.6% decrease in *Nr3c1* mRNA expression in the CeA compared to *GR*^*CRF-Ctrl*^ mice (unpaired t-test, t_10_ = 11.95, *** p < 0.001, n = 4 *GR*^*CRF-Ctrl*^ and 4 *GR*^*CRF-cKO*^ mice). Magnification 20x. Scale bar, 50 μm. (**c**) Juvenile and adult *GR*^*CRF-Ctrl*^ and *GR*^*CRF-cKO*^ mice showed similar body weight (7 weeks old: *GR*^*CRF-Ctrl*^ = 21.70 ± 0.28 g *vs. GR*^*CRF-cKO*^ = 21.78 ± 0.47 g, unpaired t-test t_14_ = 0.0977, p = 0.924; 12 weeks old: *GR*^*CRF-Ctrl*^ = 24.40 ± 0.4 g *vs. GR*^*CRF-cKO*^ = 23.4 ± 0.42 g, unpaired t-test, t_21_ = 1.654, p = 0.11). n = 4 control and 12 *GR*^*CRF-cKO*^ mice of 7 weeks old and 11 *GR*^*CRF-Ctrl*^ and 15 *GR*^*CRF-cKO*^ mice of 12 weeks old. (**d**) *GR*^*CRF-cKO*^ mice showed a slight but significant increase in *Nr3c1* expression level in the pituitary compared to *GR*^*CRF-Ctrl*^ mice (right bar graph: 24.7 ± 7.73 % increase, unpaired t-test, t_17_ = 2.156, p = 0.045), while no difference was observed in the hippocampus (left bar graph: 93 ± 5.3 % of *GR*^*CRF-Ctrl*^, unpaired t-test, t_18_ = 0.94, p = 0.358).

**Supplementary figure 4:**
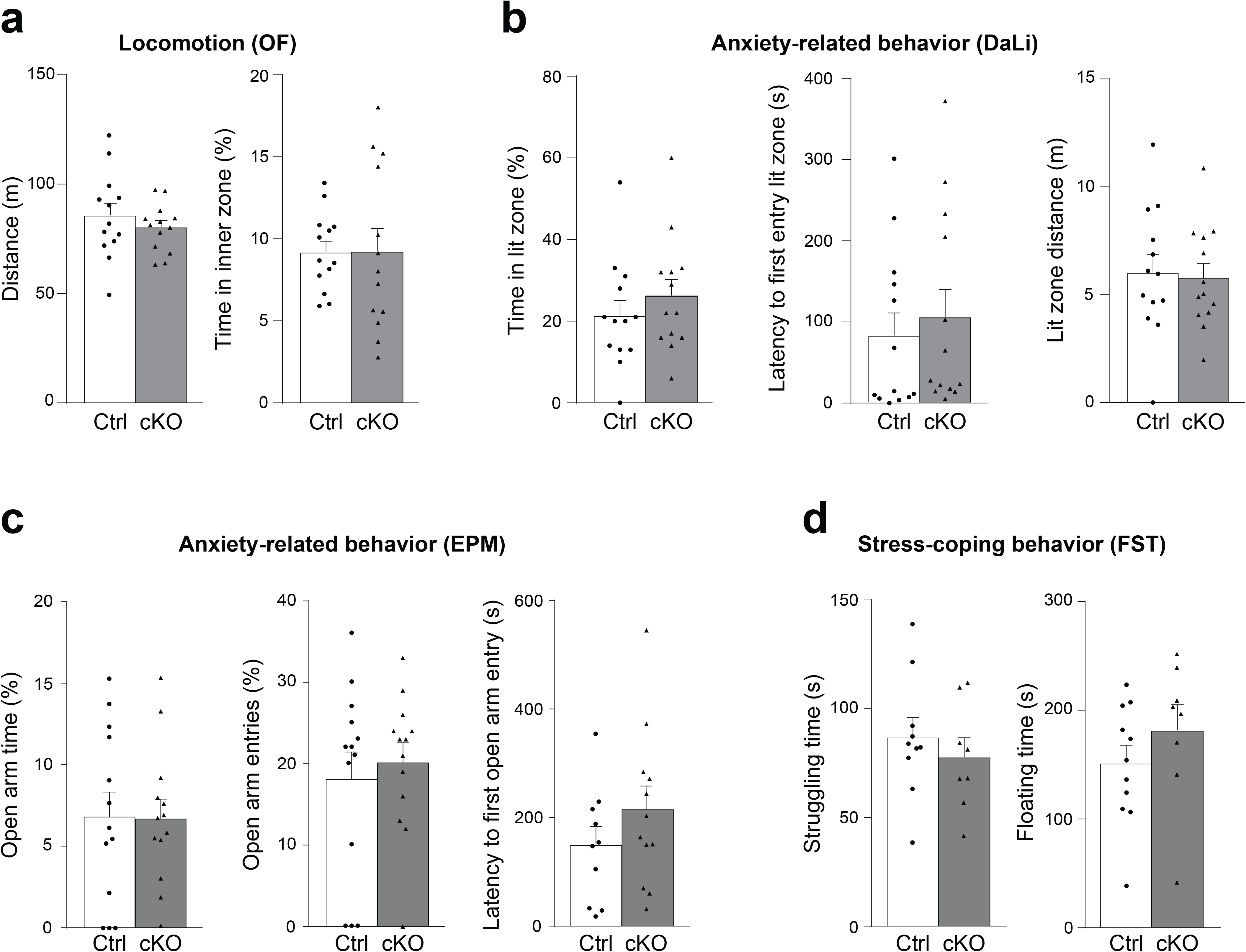
Behavioral characterization of *GR*^*CRF-cKO*^ mice. (**a**) GR deletion in CRF expressing neurons does not affect locomotor activity measured in the open field (OF: Distance travelled: unpaired t-test, t_24_ = 0.874, p = 0.39; % Time in inner zone: unpaired t-test, t_24_ = 0.03, p = 0.97). (**b**) GR deletion in CRF expressing neurons does not affect anxiety-related behavior in the dark - light box (DaLi: % Time in lit zone: unpaired t-test, t_24_ = 0.91, p = 0.37; Latency to 1^st^ entry in lit zone: unpaired t-test, t_24_ = 0.51, p = 0.61; Distance travelled in lit zone, t_24_ = 0.23, p = 0.81). (**c**) GR deletion in CRF expressing neurons does not affect anxiety-related behavior in the elevated plus maze (EPM: % Time on the open arm: unpaired t-test, t_24_ = 0.06, p = 0.95; % Entries on the open arm: unpaired t-test, t_24_ = 0.51, p = 0.61; Latency to 1^st^ entry on open arm: unpaired t-test, t_20_ = 1.19, p = 0.24). (**d**) GR deletion in CRF expressing neurons does not affect stress-coping behavior in the forced-swim test (FST: Struggling time: unpaired t-test, t_16_ = 0.71, p = 0.48; Floating time: unpaired t-test, t_16_ = 1.08, p = 0.29).

**Supplementary figure 5:**
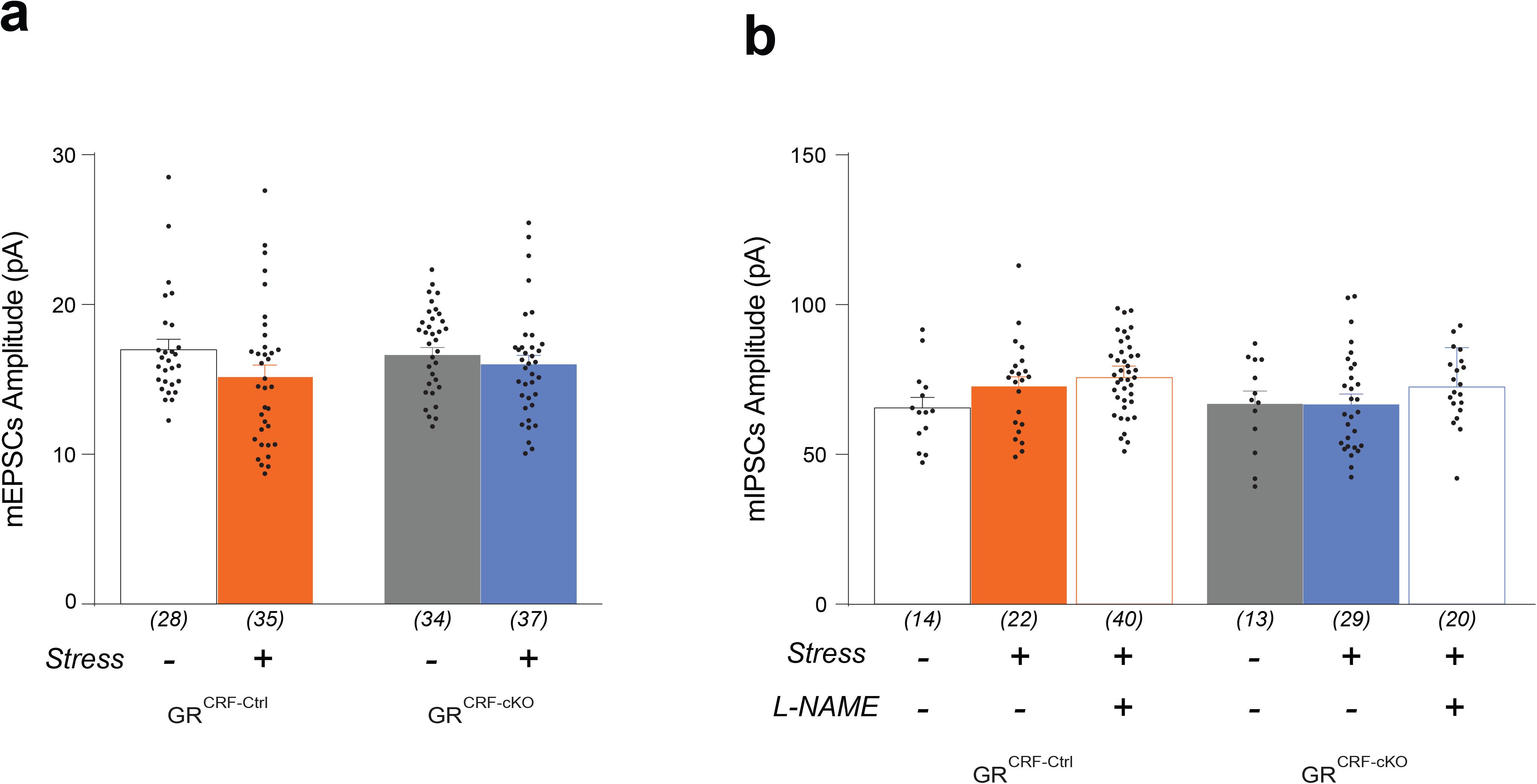
The amplitude of miniature excitatory post-synaptic currents (mEPSCs) and miniature inhibitory post-synaptic currents (mIPSCs) are not affected by GR deletion nor by repeated restraint stress (RRS). (**a**) The amplitude of mEPSCs is not affected by GR deletion, nor by RRS (2-way ANOVA Stress*Genotype F_(1,126)_ = 0.922, p = 0.338: *GR*^*CRF-Ctrl*^ basal = 16.99 ± 0.67 pA; *GR*^*CRF-Ctrl*^ RRS = 15.16 ± 0.79 pA; *GR*^*CRF-cKO*^ basal = 16.57 ± 0.45 pA; *GR*^*CRF-cKO*^ RRS = 16.00 ± 0.59 pA). (**b**) The amplitude of mIPSCs is not affected by GR deletion, nor by RRS (2-way ANOVA Stress*Genotype F_(1,70)_ = 0.902, p = 0345: *GR*^*CRF-Ctrl*^ basal = 65.55 ± 3.5 pA; *GR*^*CRF-Ctrl*^ RRS = 72.59 ± 3.28 pA; *GR*^*CRF-cKO*^ basal = 66.87 ± 4.32 pA; *GR*^*CRF-cKO*^ _RRS_ = 66.70 ± 3.45 pA). Moreover, slice incubation in the nitric oxide synthase 1 inhibitor L-NAME also has no effect on mIPSCs amplitude (*GR*^*CRF-Ctrl*^ RRS = 72.59 ± 3.28 pA *vs. GR*^*CRF-Ctrl*^ RRS + L-NAME = 75.73 ± 2.25 pA, unpaired t-test, t_60_ = 0.81, p = 0.417; and *GR*^*CRF-cKO*^ RRS = 66.70 ± 3.45 pA *vs. GR*^*CRF-cKO*^ RRS + L-NAME = 72.60 ± 3.16 pA, unpaired t-test, t_47_ = 1.194, p = 0.239). Cell numbers are indicated in brackets under each bar and were obtained from at least 7 mice.

## Online methods

### Animals and animal care

All experimental protocols were approved by the Ethics Committee for the Care and Use of Laboratory Animals of the government of Upper Bavaria, Germany. Mice were bred in the animal facility of the Max Planck Institute of Biochemistry (Martinsried, Germany) and group housed (4 to 5 mice per cage) until two weeks before the start of the experiments, when mice were single housed. Mice were housed in IVC cages, according to institutional guidelines, under a pathogen-free, temperature-controlled (23 ± 1°C) and constant humidity (55 ± 10%) on a 12-h light-dark cycle (lights on at 7 a.m.) with food and water provided *ad libitum*, at the Max Planck Institute of Psychiatry (Munich, Germany). All experiments were performed in 8- to 12-week-old (at the beginning of the experiment) male mice, during the light cycle.

Colocalization of CRF-expressing neurons and GR was carried out in *Crf*^*tdTomato*^ reporter mice resulting from the breeding of *Crf-ires-Cre* line [strain B6(Cg)-Crh^tm1(cre)Zjh/J10^, JAX stock no: 012704] and *Ai9* (*R26*^*CAG::loxP-STOP-loxP-tdTomato*^, JAX stock no: 007905). Conditional transgenic mice lacking GR in CRF-expressing neurons (GR^CRF-cKO^) were generated by crossing *Crf-ires-Cre* with *GR*^*floxed*^ mice [B6.129P2-Nr3c1^tm2Gsc/Ieg11^, EMMA strain #02124]. *Crf-ires-Cre* and *GR*^*floxed*^ mice were a generous gift from Dr. Josh Huang (Cold Spring Harbor, USA) and Dr. Günther Schütz (DKFZ, German Cancer Research Center, Heidelberg, Germany) respectively.

### Genotyping of conditional GR knockout mice (GR^CRF-cKO^)

Genotyping was performed by PCR using the following primers to identify the floxed GR allele: GR-flox1, 5 ́ GGC ATG CAC ATT ACG GCC TTC T 3 ́; GR-flox8, 5 ́ CCT TCT CAT TCC ATG TCA GCA TGT 3 ́ and GR-flox4, 5 ́ GTG TAG CAG CCA GCT TAC AGG A 3 ́. Standard PCR conditions resulted in a 225-bp wild-type and a 275-bp floxed PCR product. A premature deletion of the floxed allele would have been identified by the occurrence of a 390-bp PCR product. The following primers were used to identify *Crf-ires-Cre* mutants: 5′ CTT ACA CAT TTC GTC CTA GCC 3’ and 5′ CAA TGT ATC TTA TCA TGT CTG GAT CC 3′ (468 base pair resultant PCR band).

### *In situ* hybridization (ISH)

Single ISH procedures were performed as previously described^15,16^. Mice were killed in the morning (~ 9 a.m.) by an overdose of isoflurane. Brains were carefully removed and immediately snap-frozen on dry ice. Frozen brains were cut on a cryostat in 20-μm thick sections and mounted on SuperFrost^®^ Plus slides. The following riboprobes were used: *Crh* (3′ UTR): bp 2108-2370 bp of AY128673, *Tomato*: bp 740-1428 of AY678269, *Nr3c1*: bp 1267-1387 of NM_0081173.3 (exon 3), *c-fos*: bp 608-978 of NM_010234.

Specific riboprobes were generated by PCR applying T7 and T3 or SP6 primers using plasmids containing the above-mentioned cDNAs as templates. Radiolabeled sense and antisense cRNA probes were generated from the respective PCR products by *in vitro* transcription with 35S-UTP using T7 and T3 or SP6 RNA polymerase. Hybridization was performed overnight with a probe concentration of 7 × 106 c.p.m. ml^−1^ at 57°C and slides were washed at 64°C in 0.1x saline sodium citrate (SSC) and 0.1 M dithiothreitol. Hybridized slides were dipped in autoradiographic emulsion (type NTB2), developed after 3-6 weeks and counterstained with cresyl violet. Dark-field photomicrographs were captured with digital cameras adapted to an imaging microscope and a stereomicroscope. Images were digitalized using Axio Vision 4.5, and afterwards photomicrographs were integrated into plates using image-editing software. Only sharpness, brightness and contrast were adjusted. For an adequate comparative analysis in corresponding mutant and wild-type sections the same adjustments were undertaken. Brain slices were digitally cut out and set onto an artificial black or white background. Quantifications of ISH were performed blindly using Image J software (https://imagej.nih.gov/ij/).

### Immunofluorescence and immunoperoxidase staining

Mice were killed with an overdose of isoflurane (Floren®, Abbott) and transcardially perfused with a peristaltic pump for 3 min with ice-cold PBS containing 0.025% heparin PBS, then 5 min with ice-cold 4% PFA (w/v) in PBS, pH 7.4, and finally 1 min with PBS at a flow rate of 10 ml/min. The brains were removed and post-fixed overnight in 4% PFA at 4°C on a rotating wheel and subsequently cryoprotected in 20% (w/v) sucrose in PBS, pH 7.6 for 2-3 days at 4°C, until the brains sink to the bottom of the tube. Once saturated in sucrose, brains were snap-frozen on dry ice and stored at −80°C until sectioning (40 μm) on a sledge microtome (Leica) or freshly sectioned as follow: the day before each experiment, brains were washed with PBS and embedded in warm 6% (w/v) agarose (Invitrogen) in PBS for vibratome sectioning (HM 650V, ThermoScientific). 40 μm thick sections were prepared and stored in cryopreservation solution (25% (v/v) glycerol, 25% (v/v) ethylene glycol, 50% (v/v) PBS, pH 7.4) until free-floating IHC was performed.

For immunofluorescence staining, brain slices were washed 3 times in PBS and permeabilized with PBS containing 0.1% Triton X-100, blocked at room temperature for 1 h in PBS containing 5% normal goat serum and 0.1% Triton X-100, and incubated overnight at 4°C with the primary antibody. On the next day, slices were washed in PBS and incubated with the secondary antibody for 2 h at room temperature. After a final wash, brain slices were stained with DAPI and mounted with anti-fading fluorescence VectaShield medium (Vector Laboratories).

For immunoperoxidase staining, slices were incubated for 5 min in PBS containing 10% methanol and 3% H_2_O_2_ to quench endogenous peroxidase and then permeabilized with PBS containing 0.25% BSA, 0.5% Triton X-100 and 2% normal goat serum for 15 min at room temperature. Slices were then incubated in primary antibody overnight at room temperature, washed 3 times in PBS and incubated in secondary antibody at room temperature for 2 h. Immunoreactivity was visualized using nickel-DAB (ABC Kit and Ni-DAB kit, Vector Laboratories).

#### Antibodies

Primary antibody anti-mGR (M-20, rabbit polyclonal IgG, 1/1000, Santa Cruz Biotechnology). Secondary antibodies: Alexa Fluor 488 goat anti-rabbit IgG (1/2000, #A11034, Invitrogen Life Technologies), biotinylated goat anti-rabbit IgG (1/400, BA-1000, Vector Laboratories).

#### Image acquisition

Images were captured with either a Zeiss Axioplan2 microscope and Axio Vision 4.5 software or an Olympus IX81 inverted laser scanning confocal microscope and Fluoview 1000 Software. For confocal imaging, a z-stack of pictures of regions of interest was obtained with 0.4-1.2 μm step size and 800 × 800 to 1,024 × 1,024 pixels picture size. Images were analyzed with Image J and Imaris software (Oxford Instruments, Abingdon, UK) for colocalization experiments.

### Classical and Repeated Restraint Stress (RRS) protocol & Plasma CORT measurements

Two weeks before the experiments, male mice were separated and single housed with a 12:12 h light:dark schedule (lights on at 07:00 a.m.). To determine the basal hormone plasma levels, mice were left undisturbed throughout the night before the experiment. Blood sampling was performed in the early morning (~ 9 a.m.) and evening (~ 9 p.m.) by collecting blood from tail cut, with the time from first handling of the animal to completion of bleeding not exceeding 45 sec. For evaluation of the endocrine response to stress, blood samples were collected immediately after 15 min of restraint stress, for which animals were placed in a 50 ml conical tube. A last blood sample was collected 90 min after the end of the restraint to assess the recovery period. All plasma samples were stored at −20°C until the CORT measurement. Stress experiments were performed in the morning (~ 9 a.m.). To study the habituation of the HPA axis to a repeated homotypic stressor, this restraint stress protocol (except for the recovery sample, which was omitted) was repeated 3 times with one-week interval between each stress session (RRS), where the animals were left undisturbed.

All blood samples were kept on ice until centrifuged (4°C) and plasma was removed for measurement of CORT using radioimmunoassay, according to the manufacturer’s protocol (MP Biomedicals, Eschwege, Germany)^17^. All samples were measured in duplicates and the intra- and inter-assay coefficients of variation were both below 10%.

### Dex / CRF test

In the combined Dex/CRF test, CORT secretion in GR^CRF-Ctrl^ and GR^CRF-cKO^ mice was monitored in response to a pharmacological suppression of adrenocortical activity with the GR agonist Dex and a subsequent stimulation with CRF^18^. Seven days before the actual test, a blood sample was taken from the tail vessel of the animals at 3 p.m. in order to get a basal reference value (“Untreated”). On the day of testing, the mice received an intraperitoneal injection of Dex at 9 a.m. (Dexa-ratiopharm, Ratiopharm GmbH, Ulm, Germany), followed by an injection of CRF at 3 p.m. (Ferring Arzneimittel GmbH, Kiel, Germany; dose: 0.15 mg/kg body weight, solution freshly prepared shortly before the injection, injection volume: 0.3 ml, solvent: sterile Ringer’s solution). Immediately before CRF injection, a tail blood sample was collected (“After Dex” value) and a second one 30 min later (“After CRF” value). Two independent Dex/CRF tests were performed using either a relatively high dose (2 mg/kg body weight) or a relatively low dose of Dex (0.05 mg/kg body weight)^18^. Samples were stored at −20°C and plasma CORT concentrations were measured as described above.

### Surgery, probe implantation and *in vivo* microdialysis procedure

Free CORT levels were assessed in the prefrontal cortex (PFC) by *in vivo* microdialysis as previously described^16,19^. Male adult mice (age 10-12 weeks) were anesthetized with isoflurane (3-4% v/v in air for induction and then 1.8-2% v/v in oxygen during surgery) and placed in a stereotaxic apparatus (TSE systems Inc., Germany) with adapted components to allow inhalation anaesthesia. The fur of the skull was cleaned with 70% ethanol and the scalp was opened with a sterile scalpel along the midline. A hole was drilled and a microdialysis probe guide cannula (MAB 4.15.IC, Microbiotech AB, Sweden) was implanted into the right PFC so that the tip was 2 mm above the targeted area. The coordinates for the PFC were determined based on the stereotactic atlas (Paxinos and Franklin, 2001): AP: +1.9 mm, ML: +0.4 mm, and DV: −2.0 mm from the bregma. The guide cannula was fixed to the skull with dental cement. To connect a liquid swivel and counterbalancing arm during the microdialysis experiments, a small peg was additionally attached to the skull. Animals were allowed to recover in the experimental square plexiglas home cage for one week. Metacam® was added to the drinking water at 0.25 mg/100 ml for three consecutive days after surgery. One day prior to the experiment, mice were shortly anesthetized with isoflurane (3-4% v/v in air, 1-1.5 min) and microdialysis probes (O.D. 0.24 mm, cuprophane membrane 20 kDa 2 mm, metal-free, CMA Microdialysis AB, Sweden) and connected to the perfusion lines which consisted of FEP tubing (I.D. 0.15 mm) and a liquid swivel, which was mounted at the counterbalancing system (Instech Laboratories, Plymouth Meeting, PA, USA). The probes were continuously perfused with artificial cerebrospinal fluid (in mM: NaCl, 145; KCl, 2.7; CaCl_2_, 1.2; MgCl_2_, 1.0; Na_2_HPO_4_, 2.0; pH 7.4) at a flow rate of 0.5 μl/min, using a microinfusion pump. Two hours before the experiment, the flow rate was increased to 1 μl/min. After the equilibration period, 20 min microdialysis fractions were collected in plastic vials positioned in a refrigerated microsampler (Univentor, Malta).

At the end of the microdialysis experiments, animals were decapitated under isoflurane anaesthesia, and brains were extracted and stored at −80°C. Brains were further sectioned with a cryostat, and sections were stained with cresyl violet for a histological verification of the probe’s localization. Only data from mice with correctly placed microdialysis probes were included in the analysis.

### Behavioral studies

Behavioral experiments were performed as previously described^20^.

#### Open Field (OF) Test

The OF test was used to characterize locomotor activity in a novel environment. Testing was performed in open field arenas (30 × 30 cm, light grey) evenly unaversive illuminated at low light conditions (30 lux) in order to minimize anxiety effects on locomotion. The distance traveled and the % time spent in the center zone (as a measure of the anxiety levels of the animals) was recorded for 30 min using the ANYmaze software (Stoelting, Wood Dale, IL).

#### Dark/Light-Box (DaLi) Test

The DaLi test was used to assess anxiety-related behavior. Each animal was placed in the dark compartment (< 5 lux) of the test apparatus, facing the bright lit compartment (700 lux). During the 5 min test, the time spent in each compartment (dark, tunnel and lit compartment), the latency until the first full entry (four paw criterion) and the distance traveled into the lit compartment were assessed by means of the ANYmaze software.

#### Elevated Plus-Maze (EPM) Test

The EPM was used to assess anxiety-related behavior. The apparatus was made of grey PVC and consisted of a plus-shaped platform with four intersecting arms, elevated 37 cm above the floor. Two opposing open (30 × 5 cm) and closed arms (30 × 5 × 15 cm) were connected by a central zone (5 × 5 cm). Animals were placed in the center of the apparatus facing the closed arm and were allowed to freely explore the maze for 5 min. Parameters of interest included % open arm time [open arm time (%) = open arm time / (open arm time + closed arm time)], % open arm entries, and latency to the open arm first entry, were analyzed with the ANYmaze software.

#### Forced Swim Test (FST)

The FST test was used to assess stress-coping behavior. Each animal was placed into a glass beaker (diameter 12 cm, height 24 cm) filled with water (temperature 25-26°C) to a height of 12 cm for a test period of 5 min. The parameters floating (immobility except small movements to keep balance), and struggling (vigorous attempts to escape) were recorded using the ANYmaze software and scored throughout the 5 min test period by a trained observer.

### Viral injections and mGR rescue experiments

#### Plasmid construction

The guide plasmid pRK5-mGR-Flag was a generous gift from T. Rein (MPI of Psychiatry, Munich, Germany), and the backbone plasmid pAAV-Ef1α-DIO-mCherry-WPRE-pA was purchased from Addgene (plasmid # 37093). The mGR-Flag sequence was subcloned in the backbone plasmid using the restriction enzyme AscI and NheI. The primer for PCR were: mGR-Flag NheI forward GCTAGCATGGACTACAAGGACGACGA and mGR-Flag AscI reverse GGCGCGCCTCATTTCTGATGAAACAGAA.

#### Production and purification of adeno-associated viruses (AAVs)

Packaging and purification of recombinant (r) AAVs (serotype 1/2) was conducted as previously described^21^, with the following modifications. A variant of the human embryonic kidney cells (HEK293T), containing the SV40 large T-antigen, was transfected with the AAV transfer plasmid and the helper plasmids at a molar ratio of 1:1:1 using 1 mg/ml linearized Polyethylenimine (PEI). Three days after transfection, the cells were harvested and lysed by undergoing 3 repetitive freeze-and-thaw cycles using a dry ice/ethanol bath and a 37°C water bath. The crude lysate was obtained after a centrifugation step. The rAAV particles were then purified using a Heparin Agarose Type I chromatography column^22^. After elution, the viral particles were washed with PBS using a 100000 MWCO Amicon Ultra Filter (Millipore; Cat.: UFC910024) and concentrated to a final volume of ~100 μl.

The rAAV titers were determined by qPCR and resulted in 3×10^11^ genomic particles per microliter for the AAV1/2-Ef1α-DIO-Flag-mGR. The AAV1/2-Ef1α-DIO-mCherry (titer of 1,4×10^12^ genomic particles per microliter) was used as a control (plasmid purchased from Addgene; plasmid 50462, donated by Bryan Roth).

#### mGR rescue experiments

Mice were anesthetized with isoflurane (Floren, Abbott), 2% v/v in O_2_ and placed in a stereotactic apparatus (TSE Systems Inc., Bad Homburg, Germany) with adapted components to allow mouse inhalation anesthesia. GR^CRF-cKO^ and GR^CRF-Ctrl^ mice were bilaterally injected with either AAV-EF1α-DIO-mGR or AAV-EF1α-DIO-mCherry into the dorso-lateral part of the PVN (300 nl) using a 33-gauge microinjection needle with a 10 μl syringe (Hamilton) coupled to an automated microinjection pump (World Precision Instruments Inc.) at 50 nl/min. Coordinates from bregma were as follows: AP: −0.75 mm, ML: +/− 0.25 mm, and DV: - 4.6 mm from the cortical surface. At the end of the infusion, needles were kept at the site for 10 min and then slowly withdrawn. Post-surgery recovery included Metacam supplementation (sub-cutaneous injection of 0.5 mg/kg body weight of a 0.1 mg/mL solution) for 3 days after surgery. Four weeks after virus injection, mice were subjected to RRS and CORT levels measured as described above. After completion of the experiments, mice were killed with an overdose of isoflurane and transcardially perfused with PBS followed by 4% PFA, and brains removed for subsequent analysis. Brains were sectioned (40 μm) using a vibratome (HM 650 V, Thermo Scientific) and accurate microinjection and mGR re-expression verified by GR immunostaining (or mCherry expression for the control virus).

### Electrophysiology

#### Animals

All mice used were generated from the Crf^tdTomato^ background to identify the CRF^+^ neurons. GR^CRF-Ctrl^ and GR^CRF-cKO^ mice were assigned randomly to Basal (unstressed) and RRS groups. Mice from the RRS groups were used for electrophysiological recordings one day after the third session of restraint stress.

#### Brain slices preparation

Mice were anesthetized with isoflurane and decapitated. All following steps were done in ice-cold cutting solution saturated with carbogen gas (95% O_2_/5% CO_2_). The cutting solution (pH 7.4, 340 mOsm) consisted of (in mM): 120 choline chloride, 3 KCl, 27 NaHCO_3_, 8 MgCl_2_, and 17 D-glucose. After decapitation, the brain was rapidly removed from the cranial cavity and 300 μm thick coronal slices containing the hypothalamus were cut using a vibratome (HM650 V, Thermo Scientific). Subsequently, slices were incubated for 30 min in carbogenated artificial cerebral spinal fluid (aCSF) at 34°C. The aCSF (pH 7.4; 310 mOsm) consisted of (in mM): 125 NaCl, 2.5 KCl, 25 NaHCO_3_, 1.25 NaH_2_PO_4_, 2 CaCl_2_, 1 MgCl_2_, and 10 D-glucose. Afterward, slices were stored at room temperature (23-25°C) for at least 30 min in carbogenated aCSF before patch-clamp recordings.

#### Patch-clamp recordings

All experiments were conducted at room temperature. In the recording chamber, slices were superfused with carbogenated aCSF (4−5 ml/min flow rate). Neurons of the PVN expressing tdTomato were visually identified by epifluorescence microscopy. After identification, the cell bodies of these neurons were visualized by infrared videomicroscopy and the gradient contrast system. Somatic whole-cell voltage-clamp recordings (>1 GΩ seal resistance, −70 mV holding potential) were performed using a SEC-10L amplifier (NPI Electronic, Tamm, Germany). The current was low-pass filtered at 1.3 kHz, digitized at 6.5 kHz, and stored with the standard software Pulse 8.31 (HEKA Elektronik, Lambrecht/Pfalz, Germany). Recordings were excluded from analysis if the series resistance changed more than 10%.

#### Current-clamp recordings

These recordings were performed to compare the intrinsic electrical properties of CRF neurons in the different experimental conditions. The patch-clamp electrodes (open-tip resistance 5-7 MΩ) were pulled from borosilicate glass capillaries (Harvard Apparatus, Kent, UK) on a DMZ-Universal puller (Zeitz-Instruments, Munich, Germany) and filled with a solution consisting of (in mM): 130 K-gluconate, 5 NaCl, 2 MgCl_2_, 5 D-Glucose, 10 HEPES, 0.5 EGTA, 2 Mg-ATP, 0.3 Na-GTP, 20 phosphocreatine, pH 7.3 with KOH, 305 mOsm (all substances were from Sigma-Aldrich, St. Louis, MO). All potentials were corrected for a liquid junction potential of 14 mV. The input resistance was calculated from steady-state voltage responses upon negative current injections (1,500 ms). Firing frequency was evaluated by positive current injections that induced mild firing (4-10 action potentials) of the neurons.

#### mEPSCs recordings

Slices were superfused with carbogenated aCSF containing TTX (1 μM), bicuculline methiodide (BIM, 10 μM), CGP 55845 (5 μM), and APV (50 μM) to block sodium voltage-dependent channels, GABA_A_, GABA_B_ and NMDA receptors respectively and isolate AMPAR-mediated mEPSCs. The patch-clamp electrodes (5-7 MΩ open-tip resistance) were filled with a solution consisting of (in mM): 125 Cesium MethaneSulfonate, 8 NaCl, 4 Mg-ATP, 20 phosphocreatine, 0.3 Na-GTP, 10 HEPES, 0.5 EGTA, and 5 QX-314 chloride (pH 7.2 adjusted with CsOH, 285 mOsm).

#### mIPSCs recordings

Slices were superfused with carbogenated aCSF containing TTX (1 μM), CGP 55845 (5 μM), NBQX (5 μM) and APV (50 μM) to block sodium voltage-dependent channels, GABA_B_, AMPA and NMDA receptors respectively and isolate GABA_A_R-mediated mIPSCs. The patch-clamp electrodes (5-7 MΩ open-tip resistance) were filled with a solution consisting of (in mM): 140 KCl, 5 NaCl, 2 Mg-ATP, 20 phosphocreatine, 0.3 Na-GTP, 10 HEPES, 0.1 EGTA, and 2 QX-314 chloride (pH 7.2 adjusted with KOH, 285 mOsm). In a subset of experiments, slices were pre-incubated in the NOS 1 inhibitor L-NAME (100 μM) for at least 15 min prior to and during the recordings.

#### Analysis

Offline analysis was performed using the Pulse Software for the intrinsic electrical properties and Mini Analysis Program (Synaptosoft Inc.) for mEPSCs and mIPSCs recordings. Miniature currents were automatically detected using templates from the software with a threshold for event detection set to 3 times the RMS.

### qRT-PCR

Quantification of messenger RNA levels of Nr3c1 in the hippocampus and pituitary was carried out using quantitative real-time PCR (qRT-PCR). Total RNA was reverse transcribed using the High-Capacity cDNA Reverse Transcription Kit (Applied Biosystems). Real-time PCR reactions were run in triplicate using the ABI QuantStudio 6 Flex Real-Time PCR System and data were collected using the QuantStudio Real-Time PCR software (Applied Biosystems). Expression levels were calculated using the standard curve, absolute quantification method. The endogenous expressed gene Rpl13 was used to normalize the data. The following primers were used:

*Nr3c1* Fwd: TGTGAGTTCTCCTCCGTCCA
*Nr3c1* Rev: GGTAATTGTGCTGTCCTTCCA
*Rpl13a* Fwd: CACTCTGGAGGAGAAACGGAAGG
*Rpl13a* Rev: GCAGGCATGAGGCAAACAGTC

### Single-cell RNA transcriptomics (scRNA-seq)

#### Tissue dissociation

Mice were anesthetized using isoflurane and perfused with cold PBS. Brains were quickly dissected, transferred to ice-cold carbogenated aCSF, and kept in the same solution during dissection. Sectioning was performed using a 0.5 mm stainless steel adult mouse brain matrix (Kent Scientific) and a Personna Double Edge Prep Razor Blade. A slice (approximately −0.58 mm bregma to −1.22 mm bregma) was obtained from each brain and the extended PVN was manually dissected under microscope guidance (M205C stereomicroscope Leica, Germany). PVN from five different mice were pooled and dissociated using the Papain dissociation system (Worthington Biochemical Corporation, NJ, USA) following the manufacturer’s instructions. All solutions were oxygenated with carbogen (95% O_2_ / 5% CO_2_). After this, the cell suspension was filtered with a 30 μm filter (Partec, Goerlitz, Germany) and kept in cold oxygenated aCSF.

#### Cell capture, library preparation and high-throughput sequencing

Cell suspensions of PVN with approximately 1,000,000 cells/μl were used. Cells were loaded onto two lanes of a 10X Genomics Chromium chip per factory recommendations. Reverse transcription and library preparation were performed using the 10X Genomics Single Cell v2.0 kit following the 10X Genomics protocol (San Francisco, CA, USA). The library molar concentration and fragment length were quantified by qPCR using KAPA Library Quant (Kapa Biosystems) and Bioanalyzer (Agilent high sensitivity DNA kit), respectively. The library was sequenced on a single lane of an Illumina HiSeq4000 system generating 100bp paired-end reads at a depth of ~340 million reads per sample.

#### Quality control and identification of cell clusters

Pre-processing of the data was done using the 10X Genomics Cell Ranger software version 2.1.1 in default mode. The 10X Genomics supplied reference data for the mm10 assembly and corresponding gene annotation was used for alignment and quantification. All further analysis was performed using SCANPY version 1.3.7^23^. A total of 5,179 cells were included after filtering gene counts (<750 and >6,000), UMI counts (>25,000) and fraction of mitochondrial counts (>0.2). Combat^24^ was used to remove chromium channel as batch effect from normalized data. The 4,000 most variable genes were subsequently used as input for Louvain cluster detection. Cell types were determined using a combination of marker genes identified from the literature and gene ontology for cell types using the web-based tool: mousebrain.org^25^.

### Statistical analysis

All results are presented as mean ± SEM and were analyzed by the commercially available GraphPad Prism 7 software (GraphPad Inc.). Statistical significance was defined as *p* < 0.05. Animals were randomly allocated into different experimental groups. Conditional knockout mice and control littermates were assigned to the experimental group based on genotype. No specific randomization method was used. Age-matched littermates were used as controls in all experiments. For behavioral analysis, experimenters were blind to experimental conditions. Injection sites and viral expression were confirmed for all animals by experimenters blind to behavioral results. Mice showing incorrect cannula placement or injection sites were excluded from analysis by experimenters blind to treatment.

## References

1. McEwen, B. S. Physiology and neurobiology of stress and adaptation: Central role of the brain. Physiol. Rev. 87, 873–904 (2007).

2. Ulrich-Lai, Y. M. & Herman, J. P. Neural regulation of endocrine and autonomic stress responses. Nat. Rev. Neurosci. 10, 397–409 (2009).

3. McCarty, R. Learning about stress: neural, endocrine and behavioral adaptations. Stress 19, 449–475 (2016).

4. Lightman, S. L. The neuroendocrinology of stress: A never ending story. J. Neuroendocrinol. 20, 880–884 (2008).

5. Raison, C. L. & Miller, A. H. When not enough is too much: The role of insufficient glucocorticoid signaling in the pathophysiology of stress-related disorders. Am. J. Psychiatry 160, 1554–1565 (2003).

6. de Kloet, C. S. et al. Assessment of HPA-axis function in posttraumatic stress disorder: Pharmacological and non-pharmacological challenge tests, a review. J. Psychiatr. Res. 40, 550–567 (2006).

7. Vale, W., Spiess, J., Rivier, C. & Rivier, J. Characterization of a 41-residue ovine hypothalamic peptide that stimulates secretion of corticotropin and β-endorphin. Science (80-.). 213, 1394–1397 (1981).

8. Rivier, C. & Vale, W. Modulation of stress-induced ACTH release by corticotropin-releasing factor, catecholamines and vasopressin. Nature 305, 325–327 (1983).

9. Deussing, J. M. & Chen, A. The corticotropin-releasing factor family: Physiology of the stress response. Physiol. Rev. 98, 2225–2286 (2018).

10. Herman, J. P., Cullinan, W. E. & Herman, J. P. 1-s2.0-S0166223696100692-main.pdf. 2236, (1997).

11. De Kloet, E. R., Vreugdenhil, E., Oitzl, M. S. & Joëls, M. Brain corticosteroid receptor balance in health and disease. Endocrine Reviews (1998). doi:10.1210/er.19.3.269

12. Groeneweg, F. L., Karst, H., de Kloet, E. R. & Joëls, M. Rapid non-genomic effects of corticosteroids and their role in the central stress response. J. Endocrinol. 209, 153–167 (2011).

13. Taniguchi, H. et al. A Resource of Cre Driver Lines for Genetic Targeting of GABAergic Neurons in Cerebral Cortex. Neuron 71, 995–1013 (2011).

14. Tronche, F. et al. Disruption of the glucocorticoid receptor gene in the nervous system results in reduced anxiety. Nat. Genet. 23, 99–103 (1999).

15. Cusulin, J. I. W., Füzesi, T., Inoue, W. & Bains, J. S. Glucocorticoid feedback uncovers retrograde opioid signaling at hypothalamic synapses. Nat. Neurosci. 16, 596–604 (2013).

16. Wamsteeker, J. I., Kuzmiski, J. B. & Bains, J. S. Repeated stress impairs endocannabinoid signaling in the paraventricular nucleus of the hypothalamus. J. Neurosci. (2010). doi:10.1523/JNEUROSCI.1046-10.2010

17. Nahar, J. et al. Rapid nongenomic glucocorticoid actions in male mouse hypothalamic neuroendocrine cells are dependent on the nuclear glucocorticoid receptor. Endocrinology 156, 2831–2842 (2015).

18. Di, S., Maxson, M. M., Franco, A. & Tasker, J. G. Glucocorticoids regulate glutamate and GABA synapse-specific retrograde transmission via divergent nongenomic signaling pathways. J. Neurosci. 29, 393–401 (2009).

19. Di, S., Malcher-Lopes, R., Halmos, K. C. & Tasker, J. G. Nongenomic glucocorticoid inhibition via endocannabinoid release in the hypothalamus: A fast feedback mechanism. J. Neurosci. 23, 4850–4857 (2003).

20. Kim, J. S., Han, S. Y. & Iremonger, K. J. Stress experience and hormone feedback tune distinct components of hypothalamic CRH neuron activity. Nat. Commun. 10, 5696 (2019).

## Supplementary references

10. Taniguchi, H. et al. A Resource of Cre Driver Lines for Genetic Targeting of GABAergic Neurons in Cerebral Cortex. Neuron 71, 995–1013 (2011).

11. Tronche, F. et al. Disruption of the glucocorticoid receptor gene in the nervous system results in reduced anxiety. Nat. Genet. 23, 99–103 (1999).

15. Lu, A. et al. Conditional mouse mutants highlight mechanisms of corticotropin-releasing hormone effects on stress-coping behavior. Mol. Psychiatry 13, 1028–1042 (2008).

16. Refojo, D. et al. Glutamatergic and dopaminergic neurons mediate anxiogenic and anxiolytic effects of CRHR1. Science (80-.). 333, 1903–1907 (2011).

17. Touma, C. et al. Mice selected for high versus low stress reactivity: A new animal model for affective disorders. Psychoneuroendocrinology 33, 839–862 (2008).

18. Touma, C. et al. FK506 binding protein 5 shapes stress responsiveness: Modulation of neuroendocrine reactivity and coping behavior. Biol. Psychiatry 70, 928–936 (2011).

19. Anderzhanova, E. & Wotjak, C. T. Brain microdialysis and its applications in experimental neurochemistry. Cell Tissue Res. 354, 27–39 (2013).

20. Dedic, N. et al. Chronic CRH depletion from GABAergic, long-range projection neurons in the extended amygdala reduces dopamine release and increases anxiety. Nat. Neurosci. 21, (2018).

21. Pilpel, N., Landeck, N., Klugmann, M., Seeburg, P. H. & Schwarz, M. K. Rapid, reproducible transduction of select forebrain regions by targeted recombinant virus injection into the neonatal mouse brain. J. Neurosci. Methods 182, 55–63 (2009).

22. Zolotukhin, S. et al. Recombinant adeno-associated virus purification using novel methods improves infectious titer and yield. Gene Ther. 6, 973–985 (1999).

23. Wolf, F. A., Angerer, P. & Theis, F. J. SCANPY: Large-scale single-cell gene expression data analysis. Genome Biol. (2018). doi:10.1186/s13059-017-1382-0

24. Johnson, W. E., Li, C. & Rabinovic, A. Adjusting batch effects in microarray expression data using empirical Bayes methods. Biostatistics 8, 118–127 (2007).

25. Zeisel, A. et al. Molecular Architecture of the Mouse Nervous System. Cell 174, 999–1014.e22 (2018).

